# Diverse microbial communities assemble on both recalcitrant and labile carbon sources

**DOI:** 10.1101/2025.08.07.669103

**Authors:** Kaumudi H. Prabhakara, Yansong Zhao, Andrew D. Farr, Paul B. Rainey

## Abstract

Microbial community assembly is shaped by the nature of available resources, with labile carbon sources such as glucose often expected to support low diversity due to rapid growth and competitive exclusion. In contrast, recalcitrant substrates like cellulose are thought to support higher diversity through slower growth and increased niche partitioning. In previous work, we showed that compost-derived microbial communities propagated on cellulose maintained high diversity over nearly a year. To determine whether such diversity is specific to recalcitrant substrates or reflects more general features of assembly, we tracked community dynamics in three environments—cellulose paper, cellulose broth, and glucose—using daily 16S rRNA profiling over eight weeks. Communities were maintained through bi-weekly serial transfers, with five replicates per treatment, yielding a high-resolution dataset of over 800 samples. Despite originating from the same inoculum, communities diverged sharply in both taxonomic and functional composition. Cellulose environments yielded stable communities enriched in specialists, while glucose environments exhibited rapid succession and dominance by generalists. Surprisingly, all environments sustained comparably high levels of taxonomic diversity. Functional inference suggested extensive cross-feeding and resource salvaging in both cases. While the mechanisms maintaining diversity on glucose remain unresolved, our results reveal distinct assembly trajectories under simple carbon regimes and provide a foundation for future mechanistic study.

## 1 Introduction

Microbial communities underpin a wide range of ecological and industrial processes, performing complex functions through networks of interacting taxa (*1–6*). A central challenge in microbial ecology is to understand how such communities assemble: the determinants of their structure, stability, and diversity (*7, 8*). Of particular interest is how distinct communities emerge from a shared inoculum when exposed to different environmental contexts (*9–11*). While synthetic communities have provided useful mechanistic insights (*12*), they often lack the ecological complexity of natural systems (*13–18*). By contrast, natural communities are rich and complex but difficult to track and manipulate (*19–23*). Propagating complex communities in controlled conditions offers a productive middle ground.

In previous work (*24*), we propagated microbial communities from garden compost on cellulose for nearly a year and observed sustained high taxonomic diversity, with over 200 genera persisting through regular (bi-weekly) serial transfer. This led us to ask whether such diversity is a function of the resource—cellulose, a complex and recalcitrant polymer—or a more general feature of microbial community dynamics. To address this, we undertook a comparative study of community assembly on cellulose and glucose, the monomeric unit of cellulose, expecting the simpler substrate to support less diverse communities dominated by few rapidly growing taxa.

The results were unexpected. While community structure and dynamics differed markedly across environments, all three—glucose, cellulose broth, and cellulose paper—supported high levels of taxonomic diversity. Cellulose-grown communities exhibited slow, stable colonization dominated by specialists, whereas glucose-grown communities showed rapid turnover, high biomass, and a shifting cast of generalists and nitrogen-fixing taxa. Functional inference pointed to cross-feeding and niche construction as important features in both cases. Here we present a detailed account of these dynamics, based on daily 16S rRNA profiling over eight weeks, with bi-weekly serial transfers and five replicates per treatment—yielding a dense time series of over 800 samples. Although the mechanisms sustaining diversity remain to be fully resolved, the data reveal distinct ecological trajectories shaped by carbon source complexity and provide a foundation for mechanistic exploration of diversity maintenance in microbial ecosystems.

## 2 Results

Microbial communities sourced from compost were propagated in minimal media containing either glucose or cellulose as the sole carbon source. Communities (composed of five technical replicates) were incubated under static conditions for 14 days, then serially transferred to fresh media. This cycle was repeated for four Rounds (Rounds 1-4, Methods). Community dynamics were tracked via daily 16S amplicon sequencing, and the Amplicon Sequence Variants (ASVs) were used for analysis. For cellulose treatments, we distinguished between cells attached to the cellulose paper (CP) and those growing planktonically in the surrounding broth (CB).

### 2.1 Rapid growth-death dynamics in glucose, slow and steady dynamics in cellulose

Community biomass, estimated using DNA concentration (Figure 1), revealed different dynamics across treatments. In CP communities, biomass gradually increased with minimal loss across Rounds, consistent with the recalcitrant nature of the carbon source, though with reduced yield in later Rounds (Figure 1A). In CB, biomass lagged behind CP but peaked late in Round 1, likely reflecting the liberation of soluble carbon by CP-associated cellulolytic taxa (Figure 1B).

**Figure 1:**
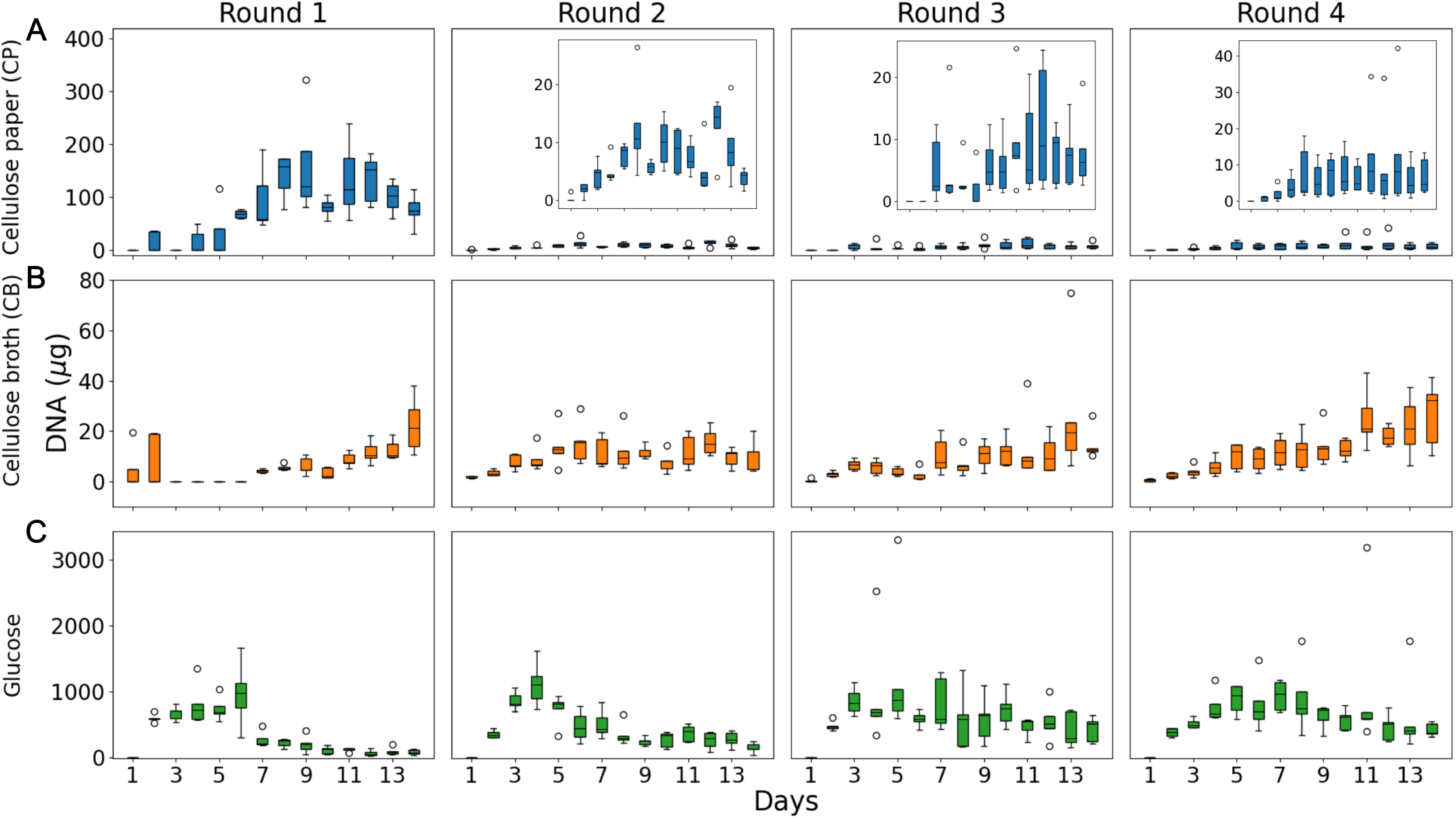
Community growth dynamics: The total DNA concentration (in *µ*g) is plotted on the y axis over the 14-day growth period (x-axis) across four sequential Rounds (columns). Each boxplot represents the values across 5 replicate communities in each environment, where the boxes extend from the first to the third quartile, solid line represents the median, and the whiskers extend from the box to the farthest data point lying within 1.5 times the inter-quartile range. **A** shows the DNA concentration for communities on cellulose paper (CP), **B** for communities in cellulose broth (CB), and **C** for communities in glucose. Insets in A show rescaled y-axes to highlight lower DNA concentrations in cellulose environments.

In glucose-supported communities biomass was 10 to 100-fold higher than in cellulose, but exhibited boom-bust dynamics. In Rounds 1 and 2, biomass peaked by day 5 before dropping sharply, consistent with rapid resource depletion and cell death. Fluctuations were more stable in Rounds 3 and 4 (Figure 1C).

To investigate these dynamics, we focused on high-abundance ASVs (those with relative abundance *>* 15%). All environments harbored transient colonizers, typically fast-growing *Pseudomonas* (Figures 2, S2–S4). In CP and CB, the taxon Cytophaga_0 dominated and remained stable over time (Figures 2A,B), mirroring biomass dynamics. In contrast, glucose environments showed rapid early expansion of dominant ASVs in each Round, followed by their replacement—closely tracking biomass turnover (Figure 2C).

**Figure 2:**
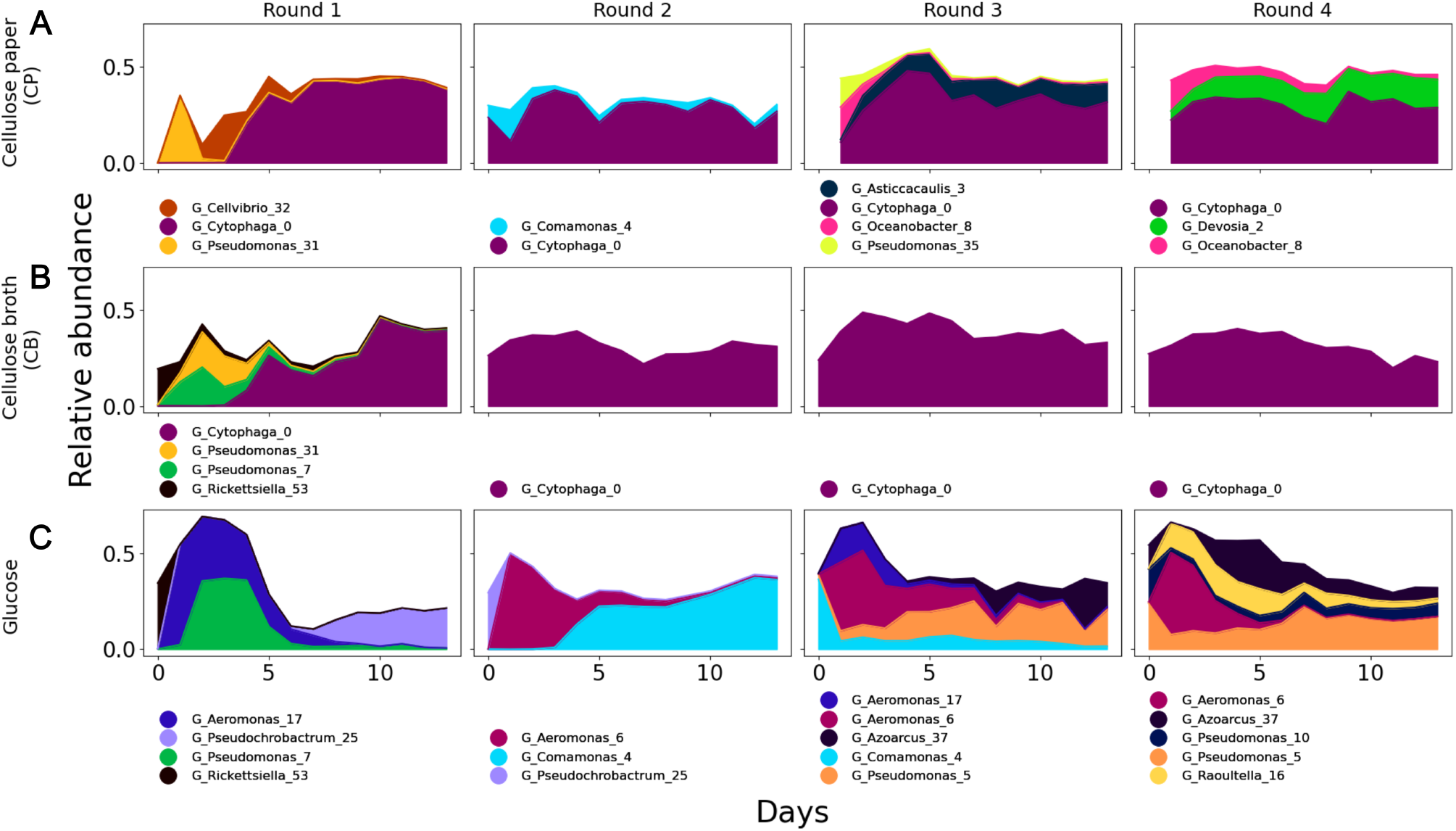
Dynamics of dominant taxa: Dominant taxa are defined as those taxa whose abundance was at least 15% in any community at any single time point (Methods). Only a single replicate for each of the three conditions across the transfers are shown. The x-axis represents days within each growth cycle, the y-axis represents relative abundance. **A** corresponds to communities on cellulose paper (CP), **B** to communities in cellulose broth (CB), and **C** to communities in glucose. Below each panel the identity of the ASVs are presented. G represents Genus, the first taxonomic classification provided by SILVA. The numbers at the end represent unique identifiers for each ASV. Figures S2-S4 show the dynamics for the other replicates.

### 2.2 Similar diversity across environments despite biomass differences

Given the differences in biomass across environments, we asked whether diversity metrics also varied. We computed Shannon diversity, richness, evenness, and the number of ASVs across relative abundance ranges for all communities (Figures 3, S5-S7). Overall, diversity metrics were broadly similar across treatments.

**Figure 3:**
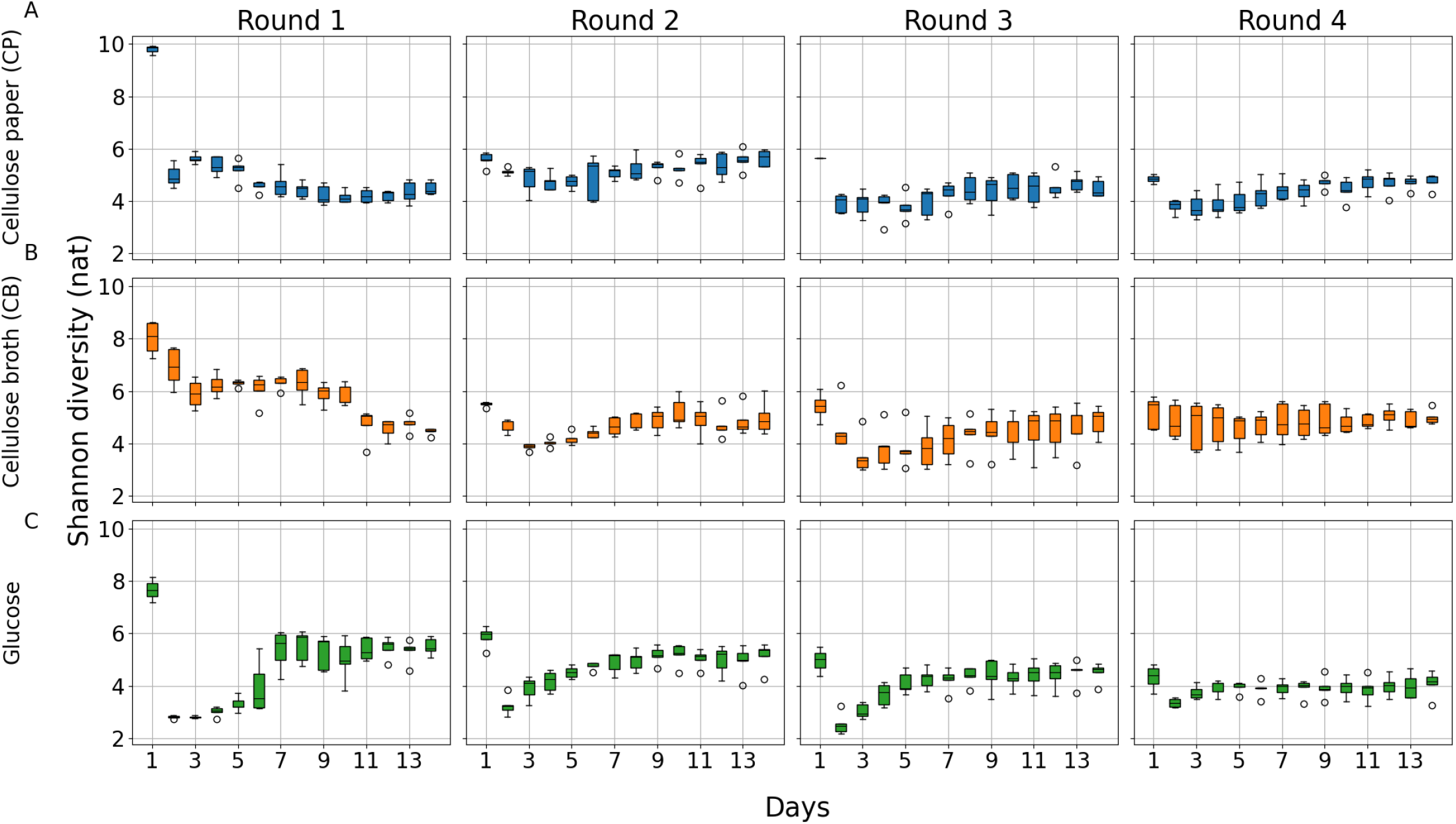
Shannon diversity dynamics across environments: Alpha diversity was quantified using the Shannon index (y-axis) and tracked over 14 days of growth (x-axis) across four Rounds of serial transfer (columns). Rows correspond to environments: cellulose broth (top), cellulose paper (middle), and glucose (bottom). Each boxplot represents values across 5 replicate communities in each environment, where the boxes extend from the first to the third quartile, solid line represents the median, and the whiskers extend from the box to the farthest data point lying within 1.5 times the inter-quartile range.

In glucose, particularly on days 2–5 of Rounds 1, 2, and 4, there were more dominant strains (with *>* 10% relative abundance) than in cellulose (Figure S5). However, at other time-points, the number of ASVs in the 5–10% and 1–5% ranges was comparable across environments, accounting for the similar overall diversity.

Shannon diversity was highest on day 1 across all environments and Rounds, followed by a sharp decline (Figure 3). This pattern was mirrored by richness (Figure S6) and evenness (Figure S7). The elevated diversity on day 1 in Round 1 reflects the high richness of the inocula, prior to acclimatization. In subsequent Rounds, transferred inocula contained ASVs that only began growing later in the Round. The early drop in diversity reflects the decline of these taxa, while the later recovery corresponds to their resurgence once appropriate substrates became available.

In glucose, evenness increased after day 5, following the initial expansion of dominant glucose consumers (Figure S7, bottom row). Prior to day 5, a few strains dominated, reducing evenness (Figure S5, bottom row). After day 5, additional taxa rose in abundance, increasing evenness (Figures 2C, S4-S5 bottom row). In contrast, evenness remained relatively stable across days in CP and CB (Figure S7, top rows), consistent with the steady dynamics discussed earlier.

To further examine diversity trends, we compared the diversity metrics averaged across days 13 and 14 of each Round, by which time communities had stabilized. By the end of Round 1, glucose communities were significantly richer, more even, and more diverse than CB communities (Figure 3, column 1; Figures S6-S7; *p <* 0.008). By the end of Round 4, however, CB communities were richer and more diverse than those in glucose (Figure 3, column 4; Figure S6; *p <* 0.05). Over time, Shannon diversity in glucose declined significantly (p = 3.8e-07), whereas it increased in CB (p = 0.026).

As expected, richness declined across Rounds in all environments due to strain loss during transfers (Figure S6). In glucose, the averaged final evenness did not change over Rounds, whereas in CB it increased significantly (p = 5.6e-06; Figure S7). In Round 1, cellulose environments were dominated by one or two cellulolytic taxa (Figures 2A,B; S2-S3, S5), leading to low evenness. In subsequent Rounds, cross-feeders (ASVs unable to directly degrade cellulose) increased in abundance, raising evenness. In glucose, cross-feeders appeared as early as Round 1 (Figures 2C, S4), resulting in stable evenness across Rounds.

Taken together, these trends explain the observed patterns in Shannon diversity: in glucose, declining richness and stable evenness reduced diversity, while in cellulose, increasing evenness countered the loss in richness, resulting in higher overall diversity.

When phylogenetic information was incorporated into diversity calculations (Figure S8), glucose communities showed a marked increase in phylogenetic diversity by the end of each Round. Initially, closely related taxa dominated, but over time more distantly related taxa—likely cross-feeders—rose in abundance. This pattern was less pronounced in the cellulose treatments.

### 2.3 Communities on cellulose and glucose remain taxonomically distinct, while replicates become similar over time

Having examined patterns in biomass and diversity, we next turned to community composition across environments and Rounds. The dynamics of dominant ASVs (Figures 2, S2-S4) clearly show that glucose- and cellulose-grown communities assembled distinct taxa, with CP and CB communities exhibiting similar compositions. This was further confirmed by Principal Component Analysis (PCA; Figure 4A-D). On day 1 of Round 1, communities clustered together (dark blue circles around coordinate (10,10) in Figures 4B-D), indicating initial similarity driven by the shared inoculum. However, as time progressed, glucose and cellulose communities diverged rapidly along the first principal component (PC1).

**Figure 4:**
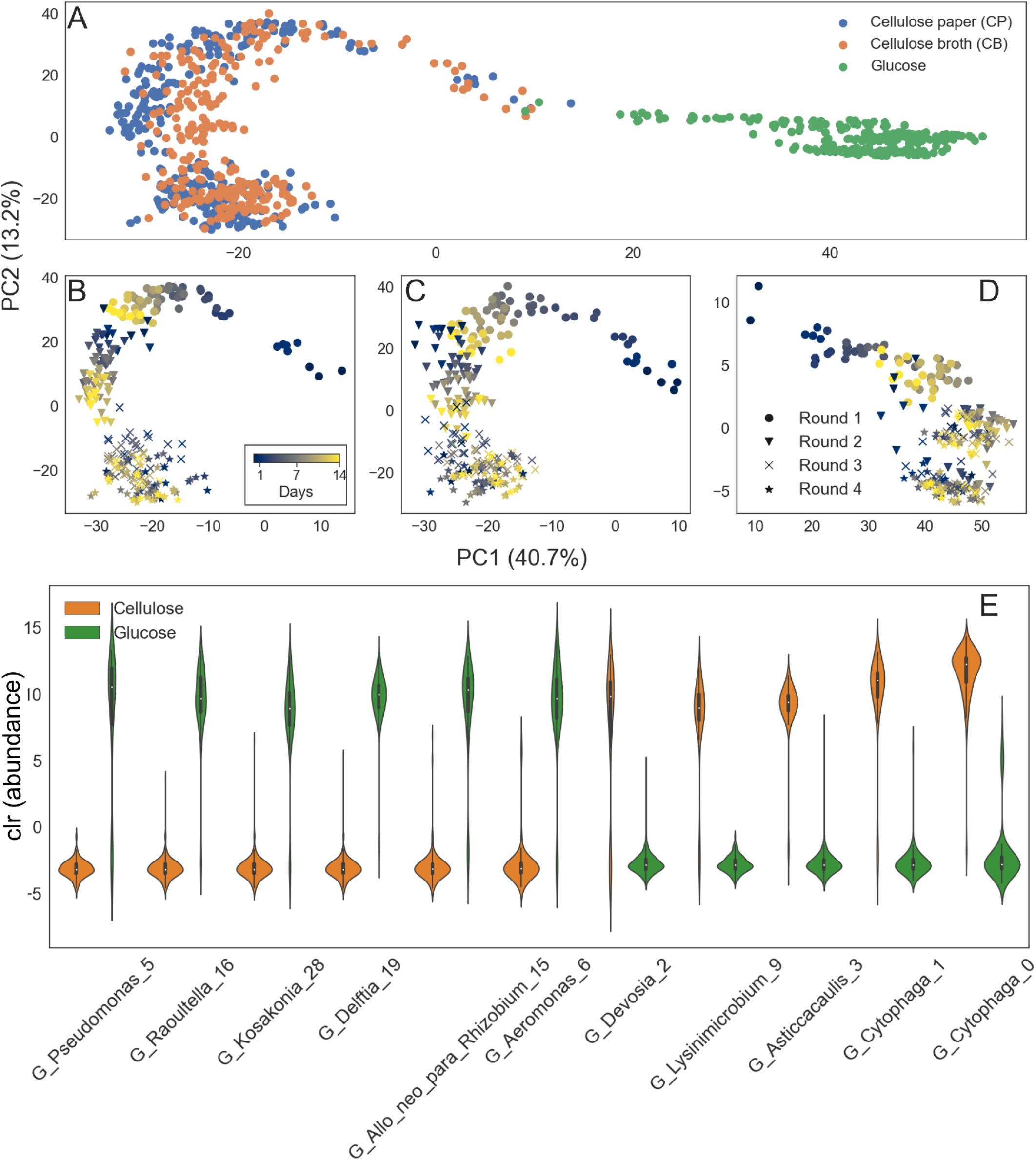
PCA on 16s sequence data: PCA was performed on the reduced dataset as described in Methods. **A**. The output of PCA plotted in two dimensions. Samples are color coded by environment. **B-D**. The same embedding as in A, but separated by environment. B shows the samples on cellulose paper (CP), C shows the samples in cellulose broth (CB) and D shows the samples in glucose. Samples are color coded by day, and markers represent the Rounds. **E**. The log transformed abundances of taxa in cellulose and glucose environments that contribute most to separation of the two environments along PC1.

Successional dynamics differed across environments. In Figure 4B–D, samples are color-coded by day and shaped by Round. In cellulose environments (CP and CB), communities showed gradual change across Rounds 1 and 2, progressing from dark blue (day 1) to yellow (day 14), and into subsequent Rounds. After Round 2, communities clustered more tightly, indicating that a steady state had been reached. In glucose, a similar pattern was observed during Round 1, but community structure stabilized by Round 2. These trends were corroborated by three beta-diversity metrics independent of PCA: Jensen-Shannon Divergence (JSD), UniFrac distance, and Aitchison distance (Figures S10, S12, S14).

Analysis of distances between technical replicates (Figures S9, S11, S13) revealed that within-Round replicate divergence decreased over time. Nonetheless, JSD and Aitchison distances showed increasing variance across Rounds, indicating that while communities generally became more similar, subgroups of replicates diverged from others. This is consistent with the emergence of dominant ASVs unique to individual replicates by Round 4 (Figures S2–S4). For example, on CP, Cytophaga_0 was a dominant member in replicates 1, 3-5, at Round 3, but a minor component of replicates 2, 3 and 5 at Round 4, with, in these replicates Oceanobacter 8 having seemingly displaced Cytophaga_0 (Figures 2A, S3). In CB, Aeromonas_6 is the principle ASV across replicates in Rounds 1-3, but Nitrosomonas_24 dominates in replicate 5 at Round 4 (Figures 2B, S2). In glucose, Pseudomonas_5 is a defining feature of later Rounds, but is at low abundance in replicate 2 at Round 4 (Figures 2C, S4). For UniFrac (Figure S13), which includes phylogenetic information, this increase in variance was not observed.

To assess the behavior of low-abundance taxa, we recalculated JSD using only ASVs with a maximum relative abundance of 1%. In contrast to full-community analyses, these comparisons (Figure S15A-C) revealed that low-abundance taxa were highly variable across replicates. Replicates clustered into two or three distinct groups with divergent compositions. Each group exhibited internal succession, but distances between groups increased over time (Figure S15G). Variance in glucose was higher than in cellulose. While dominant taxa were largely shared across replicates, low-abundance taxa varied substantially.

### 2.4 Specialists favored in cellulose environments while generalists dominate in glucose environments

We next examined the taxa that contributed to differences between communities grown on cellulose versus glucose. In cellulose environments (Figures 2A-B, S2, S3), Round 1 was characterized by transient growth of Pseudomonas_31, followed by dominance of the cellulose-degrading strain Cytophaga_0 (*25*). On CP, Cellvibrio_32 also became abundant (Figure 2A). Members of the genus *Cellvibrio* are known to produce glucosidase, a key enzyme in cellulose degradation (*26*). In subsequent Rounds, Cytophaga_0 remained dominant. Additional taxa also rose in CP communities (Figure S3), including Asticcacaulis_3 and Lysinimicrobium_9, both producers of *α*- and *β*-glucosidases (*27, 28*). Strains from the genera *Comamonas* and *Devosia*, known for metabolizing aromatic hydrocarbons (*29, 30*), also became abundant, as did marine-associated *Oceanobacter* species (*31, 32*), which consume long-chain and branched alkanes (*33*). In some replicates, *Nitrosomonas* strains appeared in later Rounds (Figures S2-S3); these ammonia-oxidizing bacteria (*34*), which fix carbon via CO_2_, may have competed with cellulose degraders for nitrogen. Overall, taxa enriched in cellulose were either direct cellulose degraders, users of alternative energy sources like ammonia, or specialists in metabolizing specific intermediates.

The dynamics of closely related ASVs offer further insight. While Cytophaga_0 was dominant across replicates and Rounds, two closely related ASVs (differing by one nucleotide) co-existed in both CP and CB (Figure S16A, D). This is unexpected given that closely related taxa are likely to compete for the same niche. In contrast, the dominant *Nitrosomonas* strains emerging by Round 4 had no closely related variants in the community (Figure S16B, C, E, F).

In glucose, early dominance in Round 1 came from *Aeromonas* and *Pseudomonas* (Figures 2C, S4). In Round 2, Aeromonas_6 became dominant and persisted through serial dilutions. In Rounds 3 and 4, additional strains of *Pseudomonas* and *Aeromonas* rose in abundance. Nitrogen-fixing taxa such as Azoarcus_37 (*35, 36*) and Allorhizobium-Neorhizobium-Pararhizobium-Rhizobium_15 became prominent in the final Rounds. Thus, glucose environments favored generalist taxa like *Pseudomonas* (*37*), *Aeromonas* (*38*), and nitrogen fixers.

Fine-scale variation among closely related ASVs further highlights these dynamics. Pseudomonas_22, which differs by one nucleotide from the initially dominant Pseudomonas_7, persisted longer into Round 2 before declining (Figure S17A). Pseudomonas_5 became dominant by Rounds 3 and 4, alongside Pseudomonas_10, which differs from it by one nucleotide (Figure S17B). Aeromonas_6, dominant in Rounds 2-4, is nearly identical to Aeromonas_17, which was dominant in Round 1 but rare thereafter (Figure S17C). In contrast, Azoarcus_37 (Figure S17D), which dominated in three replicates, had no close sequence variant.

Principal Component Analysis supports these taxonomic distinctions. PC1 separates cellulose and glucose communities (Figure 4A). Loadings along the first eigenvector (Methods) identify the taxa most responsible for this separation. Log-transformed abundances of these taxa (Figure 4E) confirm that cellulose is characterized by degraders like *Cytophaga, Lysinimicrobium, Asticcacaulis*, and metabolizers of aromatic compounds like *Devosia*, while glucose is enriched for generalists such as *Pseudomonas, Aeromonas*, and nitrogen-fixing *Allorhizobium-Neorhizobium-Pararhizobium-Rhizobium*.

From Figures 4A-D, we see that while PC1 separated the environments, the communities are displaced downwards along PC2 with time. Using the eigenvector loadings along PC2, we identified the taxa that contributed most to this temporal displacement. In cellulose environments (Figure S18, top panels), generalists such as *Pseudomonas* and *Flavobacterium* (*39*) declined sharply from Round 2 to Round 4, contributing to PC2 displacement. At the same time, specialist taxa such as the cellulose-degrading Chitinophagaceae_58 (*40–42*) and ammonia-oxidizing Nitrosomonas_14 increased in abundance by Round 4, indicating temporal selection for specialists in cellulose. Although some specialist taxa such as Cellvibrionaceae_34, Cellvibrio_80, and Devosia_64 decreased in abundance by Round 4, related strains from the same genera or families remained, suggesting competition among closely related specialist lineages. In glucose communities (Figure S18, bottom panel), some generalists declined over time (e.g., Pseudomonas_22, Pseudomonas_7) or remained at low abundance (e.g., Pseudomonas_27, Flavobacterium_36), while others such as Pseudomonas_12 increased in abundance. The nitrogen-fixing strain Azoarcus_37 also contributed to displacement along PC2, rising in abundance in some replicates by Round 4 (Figure S4). These results indicate selection for generalists and nitrogen fixers in glucose over time.

### 2.5 Interactions between taxa independent of relative abundance and environment

We next asked whether taxa in the communities exhibited signs of interaction. To explore this, we used correlations in abundance as proxies for potential interactions. The SCNIC package (*43*) was employed to infer correlation networks and define modules of co-varying ASVs (Methods) within each environment. Files S1 and S2 display the network modules for positive and negative interactions, respectively. In all environments, positive interactions between ASVs outnumbered negative ones.

Although the number of positive and negative interactions per taxon were strongly correlated (Pearson r ∼0.8, p-value ∼1× 10^*−*29^), the strengths of these interactions were not. For all taxa, the number of interactions was positively correlated with the average strength, for both positive and negative interactions (Pearson r ∼0.6, p-values < 0.0003), except for positive interactions in glucose, where no correlation was observed. In glucose, ASVs showed significantly stronger positive than negative interactions (p-value < 0.003), whereas in CB and CP, the strengths of positive and negative interactions were similar.

Comparing across environments, ASVs on CP exhibited significantly more positive interactions than those in glucose or CB (p-values < 0.007), and significantly more negative interactions than ASVs in CB (p-value < 1× 10^*−*5^). This indicates that the paper environment supported more interactions than the broth.

Modules of co-varying taxa were found to contain overlapping ASVs across environments, and the average strength of interactions within these modules was consistent across conditions. This suggests that interaction patterns were largely independent of environmental context, in particular, the main source of carbon.

We also tested whether interaction patterns were linked to taxon abundance. There was no correlation between the average relative abundance of ASVs and their number of interactions in any environment except glucose (Pearson r∼ 0.6, p-value ∼9× 10^*−*6^). Likewise, average interaction strength was not correlated with abundance in any environment except for a weak correlation in CB (Pearson r ∼0.4, p-value ∼0.02). These findings indicate that observed interactions were generally independent of ASV abundance.

### 2.6 Distinct functions are evident in different environments

To explore functional differences between communities grown on glucose and cellulose, we used PICRUSt2 (*44*) and FAPROTAX (*45*) to infer functional potential from 16S rRNA gene data. PICRUSt2 provided profiles of enzyme composition, KEGG Orthologs (KOs), and metabolic pathways, while FAPROTAX predicted broader functional capabilities. These data were analyzed by direct comparison between treatments and via PCA (Methods).

The analyses revealed clear functional distinctions between communities in the two environments (Figures 5 and 6A–D, Tables S1-S4). Further, replicate communities in CP and CB remained functionally similar over time, while replicates in the glucose-supported environment displayed temporal variation in each Round (Figures S25, S27, S29, S31), reflecting the dynamics observed in the taxonomy. Across rounds, functional similarity between the replicates did not vary (Figures S25, S27, S29, S31). In addition to the major functional differences described below, distinct sets of transporters, regulatory elements, and stress response systems were enriched in each environment.

**Figure 5:**
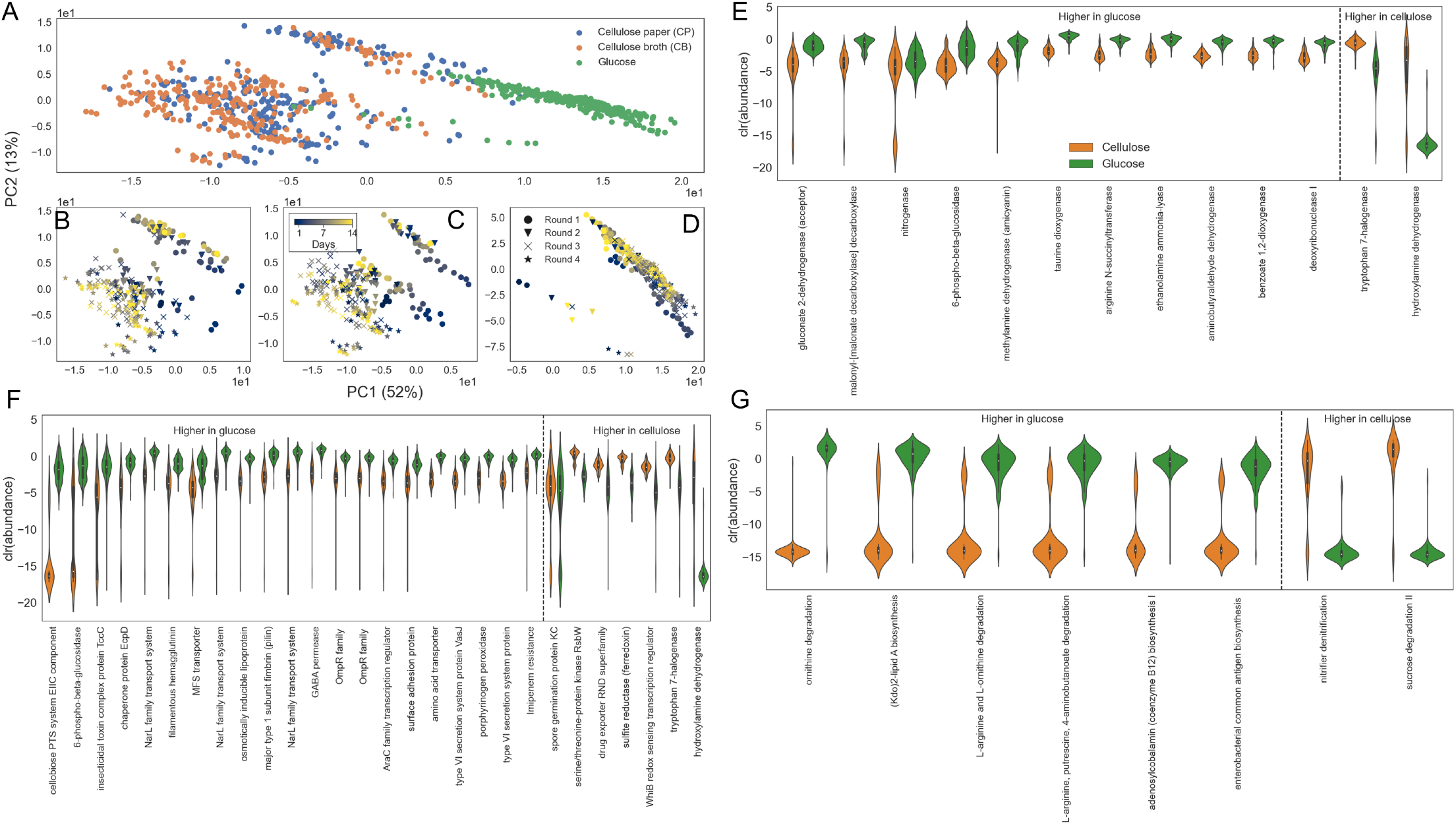
PCA on PICRUSt2 output: PCA was performed on the output of PICRUSt2 as described in Methods. **A**. Output of PCA on the abundance of EC numbers, embedded in two dimensions. The samples are color coded by their environments. **B-D**. The same embedding as in A separate by the environment. B shows the samples on cellulose paper (CP), C shows the samples in cellulose broth (CB) and D shows the samples in glucose. The samples are color coded by day, and the markers represent Rounds of transfer. **E**. The log transformed EC number number abundances in the cellulose and glucose environments contributing most to separation of the two environments is shown along PC1. **F**. The log transformed Kegg Ortholog abundances in the cellulose and glucose environments contributing the most to separation of the two environments is shown along PC1. **G**. The log transformed pathway abundances in the cellulose and glucose environments contributing the most to the separation of the two environments along PC1.

**Figure 6:**
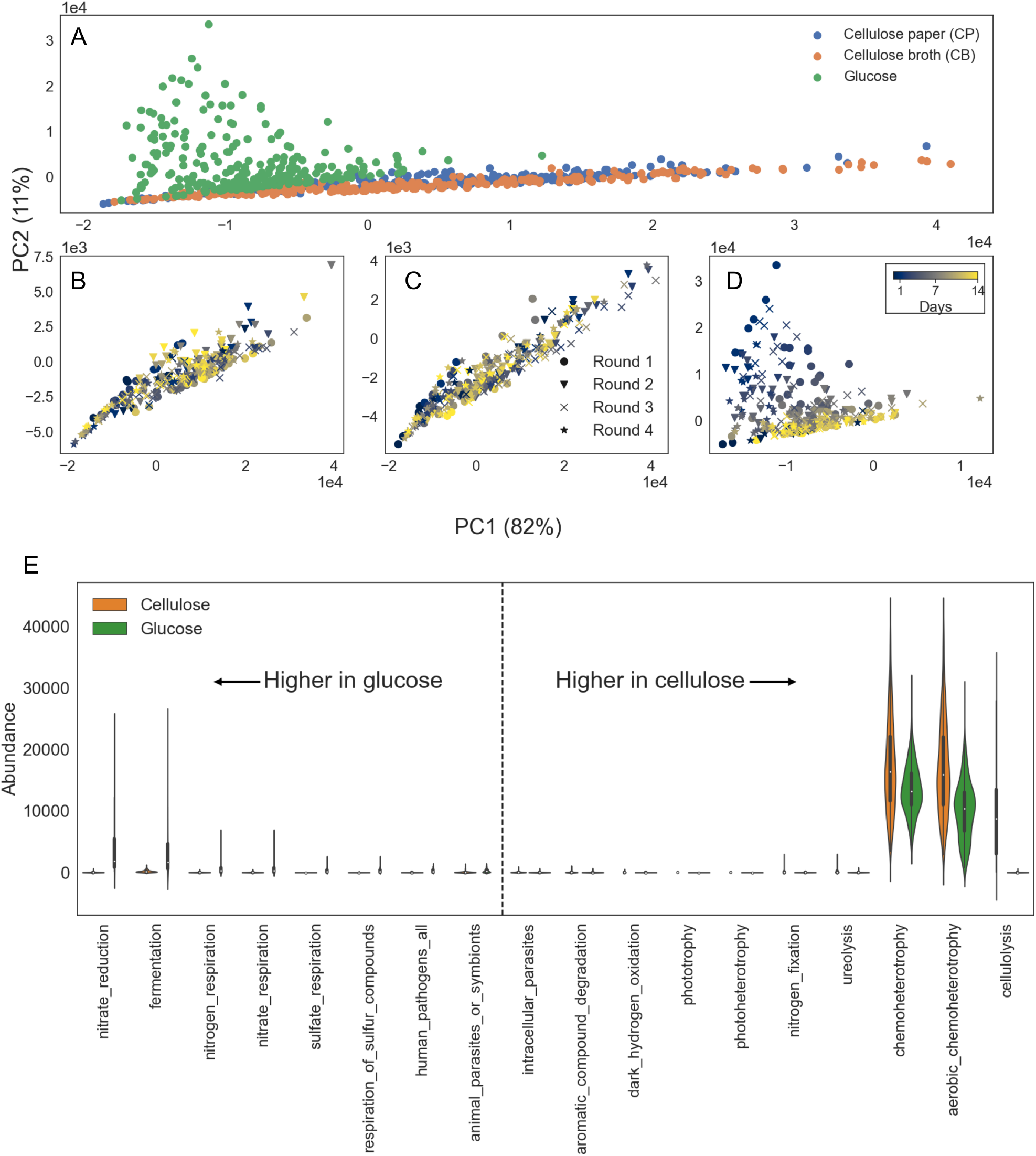
PCA on FAPROTAX output: PCA was performed on the output of FAPROTAX as described in Methods. **A**. Output of the PCA, embedded in two dimensions. The samples are color coded by environment. **B-D**. The same embedding as in A separate by the environment. B shows the samples on cellulose paper (CP), C shows the samples in cellulose broth (CB) and D shows the samples in glucose. The samples are color coded by the days, and the markers represent the Rounds. **E**. The log transformed functional number abundances in the cellulose and glucose environments contributing the most to the separation of the two environments along PC2.

#### 2.6.1 Cellulolysis, aerobic respiration, cross-feeding, and alternate energy production are prevalent in cellulose

As expected, communities grown on cellulose were enriched for genes involved in cellulose degradation (Table S4, Figure 6E), including beta-glucosidases (Tables S1 and S2). Numerous components of ABC transporter systems were also enriched (Table S2).

Aerobic respiration was more prominent in cellulose environments, evidenced by elevated abundance of enzymes related to CoA and cytochrome c (Table S1), two cytochrome c-associated KOs (Table S2), the aerobic respiration pathway (Table S3), the oxygen-dependent WhiB redox-sensing regulator (*46*) and Hydroxylamine dehydrogenase (Figure 5F).

The presence of fucosidase (Table S1), pathways for aromatic compound degradation (Figure 6E), nucleoside salvage (Table S2), and ureolysis (Table S4) suggests active cross-feeding in cellulose communities. Pathways associated with energy production independent of cellulose, such as ammonia oxidation (nitrifier denitrification, Figure 5G) and sulfate assimilation (Figure 5F), were also enriched.

Other dominant functional categories included the production and catabolism of branched-chain amino acids (Tables S1 - S3) and NAD biosynthesis (Table S3).

#### 2.6.2 Metabolic specialization, stress response, and biofilm formation are enriched in glucose communities

In the glucose environment, pathways related to the TCA cycle (Table S3) and the enzyme phosphoglycolate phosphatase (Table S1), which dephosphorylates a glycolytic intermediate, were enriched—a result consistent with glucose abundance.

High community turnover in glucose was reflected by enhanced salvaging and cross-feeding pathways. These included fermentation of formic acid (Tables S1 and S2) and increased lipid metabolism, indicated by elevated levels of acetyl-CoA C-acetyltransferase, enoyl-CoA hydratase, and acyl-ACP dehydrogenase (Tables S1 and S2). Enhanced synthesis and salvage of diverse lipids (Table S3, Figure 5G) suggest both scavenging from lysed cells and lipid storage under high-carbon conditions. This is supported by the increased degradation of xanthine, taurine, putrescine, sarcosine, and protocatechuate (Tables S1-S3, Figure 5E).

Amino acid salvage also appeared enhanced, as shown by the enrichment of polar and non-polar amino acid transporters and associated cleavage systems (Tables S1 and S2, Figure 5F). Sensory and regulatory proteins, including chemotaxis proteins, transmembrane sensors, and outer membrane receptors, were also enriched (Table S2), suggesting an increased responsive-ness to diverse metabolites. Additionally, the osmolarity-inducible lipoprotein and the MFS transporter were enriched in glucose (Figure 5F), likely reflecting adaptation to solute release following frequent cell lysis.

Functions related to nitrogen cycling were also enriched. For instance, ornithine degradation (Figure 5G), part of the urea cycle (*47*), was more abundant. Other enriched nitrogen-related functions included amidases, ureases, hippurate hydrolases (Table S2), nitronate monooxygenase involved in detoxification of alkyl nitronates (*48*) (Table S1), and multiple nitrogen respiration pathways such as nitrate reduction, nitrogen respiration, and nitrate respiration (Table S3). The NarL family transcriptional regulators (Figure 5F) and glutamine synthetase, a key enzyme for ammonium assimilation (Table S1), were also enriched.

Biofilm-related functions were a major feature of glucose communities. Components of the Type VI secretion system (Table S1; Figure 5F) (*49*), the major subunit of type 1 fimbriae (pilin), surface adhesion proteins, and filamentous hemagglutinin were all enriched (Figure 5F). These findings are consistent with the direct observation of biofilm formation in glucose cultures.

Additional functions enriched in glucose communities included the synthesis of vitamins such as pyridoxal 5-phosphate and adenosylcobalamin, the conversion of hydrogen peroxide to water, and detoxification of xenobiotic compounds.

#### 2.6.2 Distinct functions activated in the initial and final days in glucose

Figures 5D and 6D show that in the glucose environment, communities at equivalent time-points across different Rounds clustered closely. This was true for the other outputs of PICRUSt2 (Figures S22, S23) and further confirmed independently of PCA by embedding a distance metric (Figures S24, S26, S28, S32 and S33). To understand the basis of these patterns, we compared the functional profiles of communities from days 3 and 4 with those from days 13 and 14, across all Rounds (Methods). Dominant functions at each time-point are presented in Tables S5-S8. In addition to major metabolic differences, distinct stress response and resistance mechanisms were active at the two stages.

As expected, enzymes associated with sugar catabolism and TCA cycle intermediates were enriched during the early phase, including 6-phospho*-β*-glucosidase, malate dehydrogenase, and sugar phosphatases (Tables S5, S6). Interestingly, both anaerobic (e.g., formate C - acetyltransferase, pyruvate-ferredoxin flavodoxin oxidoreductase, and C4-dicarboxylate transporter (DcuC family), fermentation) and aerobic respiration pathways (e.g., pyruvate synthase) were detected early. By contrast, aerobic respiration dominated in later time-points, with enrichment of cytochrome c oxidase and related pathways (Tables S5-S7).

Salvage pathways—including amino acid and aromatic compound degradation—were already active by days 3 and 4 (Table S7), suggesting substantial early cell turnover. By the final days, communities showed elevated activity in ABC transporters, fatty acid synthesis, and lipid metabolism, consistent with storage and scavenging (Tables S5-S7).

An increase in serine transporter abundance during the early phase is noteworthy (Table S6). This transporter is known to prevent cell lysis under glucose-depleted conditions (*50*),potentially explaining how initially dominant ASVs persist across Rounds.

Nitrogen metabolism also shifted over time. While nitrate reduction was enriched in the early days, a broader suite of nitrogen-related functions—including ammonia assimilation, nitrate and nitrogen respiration, and ureolysis—was upregulated in later stages (Table S8), suggesting intensifying nitrogen competition.

In the final days, enrichment of receptors such as histidine kinases, chemotaxis receptors, and outer membrane receptors (Tables S5, S6) indicated increased sensing of and response to available metabolites, consistent with enhanced cross-feeding. Sulfate and sulfur respiration also increased (Table S8), further supporting the view of intensified resource competition over time.

## 3 Discussion

A central finding of this study is that high microbial diversity can be maintained on both recalcitrant and labile carbon sources, despite large differences in community composition, function, and biomass. From a classical ecological perspective, this outcome is surprising. Foundational theory—supported by early microbial evolution experiments in well-mixed chemostats—predicts that single limiting resources should support only a few dominant taxa due to competitive exclusion (e.g., (*51–55*)). The “paradox of the plankton” and other works (*56–58*) articulated this puzzle more broadly: how can such high diversity persist in seemingly uniform environments? More recent work, particularly in microbial systems, has begun to reconcile this paradox. Studies have shown (*9, 11, 59–68*) that metabolic cross-feeding, temporal niche generation, and eco-evolutionary feedbacks can sustain complex communities even on single carbon sources. Our results contribute to this growing body of evidence by demonstrating that distinct but comparably diverse microbial communities can assemble and persist on either glucose or cellulose—substrates that differ dramatically in structure and accessibility.

Glucose and cellulose gave rise to taxonomically and functionally distinct communities, consistent with expectations based on carbon chemistry. Glucose-supported communities exhibited rapid growth, high biomass, and early dominance by fast-growing generalists. Correspondingly, anabolic pathways were enhanced in these communities, a result expected from previous analyses of proteome allocation (*69, 70*). Cellulose-supported communities, by contrast, assembled slowly, maintained lower biomass, and were enriched for taxa known to specialize on complex polysaccharides. These differences likely reflect underlying metabolic trade-offs. As noted by (*71*), microbial investment in glycosidic enzymes—essential for cellulose degradation—is anti-correlated with investment in cell replication machinery. Such a trade-off could explain the lower overall productivity in cellulose environments. However, despite slower growth and lower resource availability, cellulose-grown communities ultimately achieved similar levels of taxonomic diversity. In fact, the cellulose-supported communities eventually became richer than the glucose-supported communities, consistent with expectations from the number of secondary metabolites produced by the two substrates (*60*). This suggests that recalcitrant substrates, while energetically costly to exploit, promote sustained diversity by enabling niche partitioning and slowing competitive exclusion.

A second explanation of the difference lies in the emergence of cross-feeding and metabolic interdependence. Functional inference revealed signatures of extensive but distinct metabolite exchange in both environments, including enrichment of chemotaxis and transporter systems, particularly in glucose-grown communities. Taxon co-occurrence patterns further indicate that low-abundance taxa are strongly connected within the community and may rely on metabolic byproducts produced by dominant taxa. This aligns with previous findings that even single-resource environments can give rise to metabolically structured communities through niche construction and resource diversification (*9, 60*). The persistence of cross-feeders at low abundance suggests that these taxa may be crucial to sustaining high overall diversity, and that ecosystem function emerges from distributed, cooperative interactions.

Third, nutrient stoichiometry and dynamic resource limitation likely shaped community composition in distinct ways. Although the growth media had similar C:N:P ratios in both conditions, the rate and mode of carbon consumption differed. In glucose, rapid uptake led to nitrogen limitation, as indicated by enrichment of nitrogen transporters and nitrogen-cycling functions, This caused strong competition, reflected in the survival of only distantly related ASVs performing nitrogen cycling functions. In cellulose, carbon was released more slowly, favoring energy-efficient respiratory strategies such as oxidative phosphorylation, reducing the intensity of nitrogen competition and promoting alternative energy production pathways. Apart from carbon, nitrogen and phosphorous, the availability of oxygen was also different - rapid growth in glucose environment likely created anaerobic pockets, while the cellulose-supported environment remained aerobic. These contrasting metabolic regimes—driven by the kinetics of substrate availability—help explain the functional divergence between communities and support the idea that nutrient imbalance can generate indirect niche dimensions even in nominally simple environments.

Despite pronounced taxonomic turnover, functional profiles remained relatively stable within each environment. Communities grown on glucose showed similar functional signatures across early time-points in different experimental Rounds, while cellulose communities maintained functional consistency throughout. This decoupling between taxonomy and function indicates strong functional redundancy—a common feature of complex microbial communities (*19, 72, 73*). In our case, it suggests that while species composition may be dynamic, the ecological roles required to process each carbon source are consistently fulfilled. Such redundancy may buffer ecosystem function against stochastic demographic fluctuations or taxon loss.

Our findings contrast with previous reports of highly reproducible assembly dynamics in glucose-supplemented communities inoculated from soil, such as those described by Goldford et al. (*9*). In that study, just 12 batch transfers (∼84 generations) in shaken culture led to the rapid emergence of simple and convergent communities dominated by a few metabolic generalists. That outcome was interpreted as evidence of universal metabolic constraints driving deterministic assembly. However, the reproducibility observed in such systems likely reflects strong ecological filtering arising from both inoculum preconditioning and the propagation regime, which favors fast-growing fermentative and respiratory taxa.

In our experiments, despite using glucose as the sole carbon source and observing initial dominance by fast-growing Pseudomonas, community dynamics were less predictable: biomass levels fluctuated sharply, dominant taxa turned over within rounds, and diversity metrics—especially evenness—varied significantly across time and replicates. Cross-feeders emerged early and maintained diversity, but selection did not drive strong convergence. These findings suggest that deterministic simplicity is not an inevitable outcome of microbial community assembly on labile carbon sources. Instead, it emerges under particular combinations of environmental structure, propagation regime, and selection intensity. Our results point to a more contingent and historically structured assembly process, even under nominally simple conditions.

Patterns of divergence across replicate communities reveal an interplay between deterministic and stochastic forces—the latter likely influential on dynamics of low abundance taxa. While dominant taxa converged within each environment—suggesting selection under shared conditions—lower-abundance ASVs diverged across replicates, especially in glucose. This divergence likely reflects increasing environmental complexity over time, driven by cell turnover, byproduct accumulation, and niche diversification. Prior work has shown that microbial communities can diverge more in complex environments (*74*), supporting our hypothesis that more diverse byproducts in glucose compared to cellulose cause stronger divergence in the low abundance taxa that depend on these byproducts.

Spatial structure further modulated community composition. In the cellulose environment, paper-associated communities differed from those in the surrounding broth. Taxa colonizing the cellulose matrix were more abundant and interacted with a greater number partners, consistent with the hypothesis that spatial partitioning can promote coexistence by reducing direct competition. While differences in diversity between compartments were modest, the presence of physical structure appears to create microenvironments that facilitate co-assembly and resource partitioning.

Although ASVs that differ by only a single nucleotide are clearly closely related, they can nevertheless differ substantially in accessory gene content. Such ASVs frequently co-occur in our communities, but it is often unclear whether they represent distinct species within the same genus or different strains of the same species. While the coexistence of multiple species within a genus is readily explained, the stable coexistence of closely related strains poses a greater challenge. One way to distinguish between these possibilities is to examine genome databases. For instance, the Genome Taxonomy Database contains only six strains assigned to the genus *Cytophaga*, but 8751 strains for *Pseudomonas*. Without further information, it remains difficult to determine whether our ASVs reflect intra- or interspecific diversity. However, this distinction is important—particularly in the context of competitive interactions (*75*)—and future work involving strain isolation and genome sequencing will be required to clarify these relationships.

Together, these findings reveal that microbial diversity can persist under far simpler resource conditions than previously assumed. Functional redundancy, emergent metabolic interactions, dynamic nutrient limitation, and spatial structure all contribute to this outcome. Rather than acting in isolation, these processes interact to shape distinct yet diverse communities across contrasting environments. Our results underscore the importance of considering not only the identity of available resources but also the ecological dynamics they generate—including feedbacks through byproduct formation, growth kinetics, and interspecies interactions. Ongoing work is aimed at disentangling these mechanisms more precisely through isolate-based reconstructions, genome-resolved metagenomics, and experimental manipulation of interaction networks. Of particular importance is understanding the relationship between diversity maintained in these experimental systems and diversity of types present in the source inoculum.

## 4 Limitations of the study

This study has two primary limitations. First, while the initial inocula derived from compost contained both prokaryotic and eukaryotic organisms, our sequencing data captures only the prokaryotic component. As a result, we may have overlooked potentially important eukaryotic contributions to community dynamics and function. Second, all functional inferences were based on 16S rRNA gene sequences, using predictive tools that rely on reference genomes. This approach assumes that taxa with similar 16S sequences share similar functional repertoires, an assumption that may not hold for diverse environmental strains. Consequently, the inferred functional profiles should be interpreted with caution.

## 5 Methods

### 5.1 Sample collection, propagation and DNA extraction

On 2nd November 2021, ∼50 g compost was sampled from a garden at 54^*°*^09^*′*^32.4”*N* and 10^*°*^26^*′*^13.2”*E*. Then, 20 g sample was mixed and vigorously shaken with 100 mL M9 solution, and then left static for ∼5 min for the solid particles to sediment. Next, 5 out of 6 microcosms with 190 mL M9 media and 40 cm^2^ cellulose paper (consisting of 9 pieces of 2 cm × 2 cm paper and 16 pieces of 0.5 cm ×0.5 cm paper), and 5 out of 6 microcosms with 190 mL M9 media and 0.2 % w/v glucose, were each inoculated with 2 mL supernatant of the compost wash. The microcosms were statically incubated in 28°C for the next 14 days. Every day during incubation, a piece of 0.5 cm × 0.5 cm paper and 2 mL broth from each cellulose microcosm, and 2 mL broth from each glucose microcosm, were harvested. The DNA in the samples were immediately extracted using Qiagen PowerSoil Pro kit, and then quantified using Qubit dsDNA High Sensitivity kit.

The quantified DNA was then scaled up to obtain the total DNA content of the communities (Figure 1). The DNA concentration was first multiplied by the volume of elution (here 50µ*L*) to get the DNA eluted. For the liquid samples, since the DNA was initially extracted from 2 mL of the culture, the eluted DNA was divided by 2 mL and multiplied by 200 mL, the volume of the culture, to estimate the DNA content of the entire culture. For the cellulose paper samples, since the DNA was harvested from 0.25 cm^2^ paper, the eluted DNA was divided by 0.25 cm^2^ and multiplied by 40 cm^2^, the total area of the paper in the culture, to estimate the total DNA content on the paper.

At the end of the 14-day incubation, 1/20 of the content of each microcosm was transferred to a new, sterile microcosm with the same resource type and amount. In detail, for each microcosm, 10 mL of the broth was used to inoculate the next microcosm; for each cellulose microcosm specifically, the 9 pieces of 2 cm × 2 cm paper were harvested and vortexed with 30 mL M9 solution into a slurry, and 1.5 mL of the slurry was used as an additional inoculum for the next microcosm. The newly-inoculated microcosms then entered the next 14-day incubation Round, with the same sampling and genomic preparation protocols as the first Round, and this repeated for a total of 4 Rounds (i.e., 56 days in total).

### 5.2 Sequencing and analysis of raw sequencing data

All the DNA samples gathered in the experiment were subject to 16S meta-barcoding for community composition determination. In detail, 16S rRNA v1-v2 regions in each sample was amplified by 30 cycles of PCR (primers 27F 5’-AGAGTTTGATCCTGGCTCAG-3’, 338R 5’-TGCTGCCTCCCGTAGGAGT-3’) according to the dual-index library preparation strategy (*76*), which were then sequenced by the Illumina MiSeq platform. Next, the R package DADA2 was used to clean the raw sequences according to (*77*), and the Silva Project’s version 138.1 prokaryotic SSU database was used for taxonomy assignment up to genus level. The raw reads, the filtered reads, and the outputs of the DADA2 pipeline are available at: https://doi.org/10.5281/zenodo.16760758.

The exact sequences, denoted as Amplicon Sequence Variants (ASVs) were used for further analyses. All analyses were performed on Python (*78, 79*) unless otherwise specified. In total, 15576 ASVs were detected in the data. Each sample had on average 24044 counts with a standard deviation of 27% (Figure S1). The highest count any ASV had in any sample was 20965.

### 5.3 Alpha diversity - diversity metrics

Diversity was calculated using three metrics for all samples without removing rare ASVs.

#### 5.3.1 Shannon diversity

The Shannon diversity metric (*80, 81*), H, is defined by Equation 1.

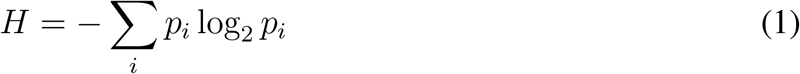

Here, *p*_*i*_ is the relative abundance of ASV_i_ in the community. This metric was calculated using skbio’s “shannon” function (*82*).

#### 5.3.2 Abundance based Coverage Estimator

The Abundance based Coverage Estimator (ACE) metric *S*_*ace*_ (*83*), measures richness, and is given by Equation 2.

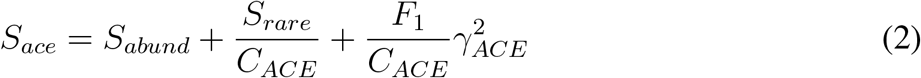

The ASVs are classified as abundant, *S*_*abund*_, and rare, *S*_*rare*_, depending on whether the their count is greater than or less than 10 respectively. *F*_*i*_ is the number of species with a count of i (i.e. *F*_1_ is the number of species with count 1). 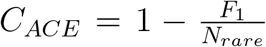 is the sample coverage, where 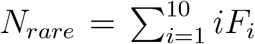 is the total number of counts in rare species. γ_*ACE*_ is the estimated coefficient of variation for the rare ASVs, given by 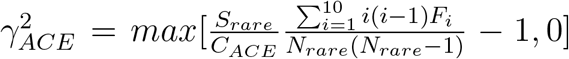. This metric was computed using skbio’s “ace” function (*82*), based on the EstimateS manual by Colwell (*84*).

#### 5.3.3 Simpson’s evenness

Simpson’s evenness measure (*85*), E, quantifies how even the samples are. It is defined by Equation 3.

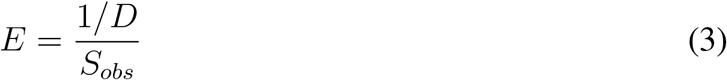

Here,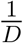 is the effective number of species, where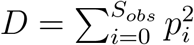. As before, *p*_*i*_ is the relative abundance of ASV_i_ in a sample. *S*_*obs*_ is the number of unique ASVs observed. This measure was calculated using skbio’s (*82*) “simpson_e” function.

#### 5.3.4 Faith’s Phylogenetic diversity

To include the phylogenetic information, we used the Faith’s Phylogenetic Diversity (PD) metric. We used scikit-bio’s (*82*) implementation of Faith’s PD metric, which is based on (*86*). The phylogenetic information is input in the form of a tree. As it is difficult to align 15,576 sequences, we reduced our data to include only those taxa that have a maximum relative abundance of at least 0.5% in any sample. This reduced the dataset to 727 ASVs. The fasta file containing the sequences of these ASVs were aligned and a tree created using SILVA’s ACT feature (*87*) with the FastTree algorithm. This tree was used as the input for skbio’s Faith’s PD calculation.

#### 5.3.5 Statistical testing

The two sample Kolomogrov-Smirnov test was used to measure significance. The null hypothesis of this test is that the two samples are drawn from the the same distribution. When a test’s p-value is low enough, the null hypothesis can be discarded. In such cases, where the null hypothesis could be discarded, the mean of the two samples was compared to know which one had the higher value. scipy’s “ks_2samp” function was used with the default options. When testing mutliple samples, the Bonferroni correction (*88*) was applied to correct the limit of significant p-value by dividing 0.05 by the number of tests.

### 5.4 Principal Component Analysis

In order to perform the Principal Component Analysis (PCA), it is necessary to take into account the compositional nature of the relative abundance data (*89, 90*).

#### 5.4.1 Zeroes and rare taxa

One of the log transformations introduced by (*91*), the center log transform (clr) was used here to overcome the limitations posed by the compositional nature of the data. The “clr” function from skbio was used to calculate the transform. However, one problem with using such log transformations is the presence of zeros in the data. This problem was overcome here by adding a small number, 0.001, to all the data. It has been shown (*92*) that adding a small pseudocounts does not impact further analyses. To further discard rare taxa, with low counts, the maximum count of each ASV across all samples was plotted (Figure S19A). This shows that there are many taxa whose maximum abundance across all samples is very low. Any ASV whose maximum count was less than 1350 was removed (vertical dashed line in Figure S19A). Given the average counts per sample, 24044, and the standard deviation is about 27% (Supplementary Figure), this implies removing ASVs whose maximum relative abundance across all samples is less than about 5.6%. With this thresholding, the number of sequences reduces from 15576 to 102.

#### 5.4.2 Choosing the number of dimensions

PCA was performed on the reduced dataset using scikit-learn (*93*). The distribution of eigenvalues is plotted in Figure S19B, where it is noted that the data had two dominant eigenvalues, while the rest were comparatively low. The first two eigenvalues explain 40.7% and 13.2% of the variance. Figure S19C shows the cumulative explained variance. These indicated that the first two dimensions explained the data well. To further confirm this, the data were shuffled by randomizing the counts for each sample. The clr transform was performed on this randomized dataset. By performing PCA on this randomized clr transformed dataset, the eigenvalue distribution of the randomized dataset was obtained (Figure S19D,E). The two dominant eigenvalues observed before were absent, indicating that the structure of the dataset is lost on randomizing.

#### 5.4.3 PCA without removal of rare taxa

PCA was performed on the complete data, without removing any rare taxa, after performing the clr transform (Figure S20). The eigenvalue distribution was similar, with the first two eigenvalues much higher than the rest, although in this case, the third eigenvalue was also significant (Figure S20E). However, the first two components only explained 17.2% and 9.1% of the variance (Figure S20F). The basic structure of data remained the same as in the PCA with rare taxa removed, both across and within the environment (Figure S20A-D), especially with regard to the separation along Principal Component 1 (PC1), and the succession dynamics. Random shuffling of the data caused it lose its structure, with the high eigenvalues dropping out (Figure S20G,H).

#### 5.4.4 Contributors to separation along PC1

The contributions of the ASVs to the separation of the communities grown on glucose broth from the communities grown on cellulose along PC1 were calculated as follows (*11*). The eigenvector along PC1 is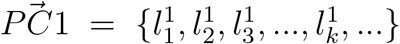, where 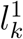 is the loading correspond-ing to ASV *k* along PC1. The superscript 1, refers to PC1. The composition of a sample is the vector, 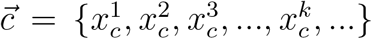, where 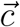 represents a sample and 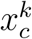 is the mean subtracted, clr transformed abundance corresponding to ASV *k* in the sample represented by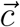. The distance between two communities along PC1 is 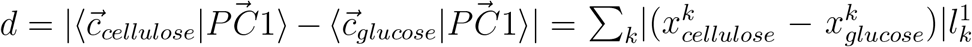, where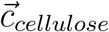 represents a community grown on cellulose and 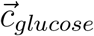 represents a community grown on glucose. The summation is over all the ASVs *k*. 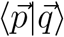denotes the inner product of the two vectors 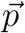 and 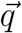 and can be interpreted as the projection of 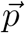 on 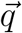. Therefore, the contribution of ASV *k* to the separation of the communities grown in glucose broth from the communities grown in cellulose along PC1 is 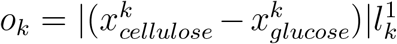.

The *o*_*k*_ for each ASV *k* was computed by considering all possible pairs of communities grown in glucose broth and cellulose broth across all replicates, transfers and days. The median across all these pairs was used as the contribution of the ASV to separation along PC1. The cumulative distribution of *o*_*k*_ was obtained for all ASVs, and a cut-off value was chosen to be at 90 percentile. The ASVs whose *o*_*k*_ was higher than this cutoff were designated as those contributing the most to the separation of the communities grown on glucose from the communities grown on cellulose along PC1. The clr transformed abundance of these ASVs are plotted in Figure 4E.

#### 5.4.5 Contributors to displacement along PC2

From Figure 4, it is clear that, the samples move down along Principal Component 2 (PC2) in all three cases. This is especially clear from Round 2 when the initial transients have ended. To know which ASVs contribute the most to this displacement along PC2, a similar method as before was applied.

The eigenvector along PC2 is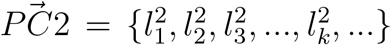, where 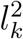 is the loading corre-sponding to ASV *k* along PC2. The superscript 2, refers to PC2. As before, the composition of a sample is the vector, 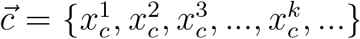, where 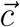 represents a sample and *x*^*k*^ is the mean subtracted, clr transformed abundance corresponding to ASV *k* in the sample represented by 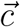. The distance between two communities along PC2 is 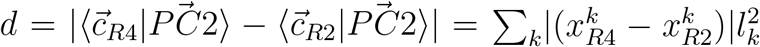, where 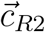 represents a community at Round 2 and 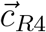 represents a com-munity at R4. The summation is over all the ASVs *k*. Therefore, the contribution of ASV *k* to the displacement of the communities across Rounds along PC2 is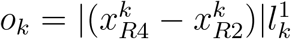.

The *o*_*k*_ for each ASV *k* was computed by considering all possible pairs of communities in Rounds 2 and 4 across all replicates, transfers and days, but within a given environment. The median across all these pairs was used as the contribution of the ASV to displacement along PC2. The cumulative distribution of *o*_*k*_ was obtained for all ASVs, and a cut-off value was chosen to be at 90 percentile. The ASVs whose *o*_*k*_ was higher than this cutoff were designated as those contributing the most to the displacement along PC2. The clr transformed abundance of these ASVs are plotted in Figure S18.

### 5.5 Beta diversity - distance metrics

Two distance metrics were computed for the data using two metrics. The entire dataset was used for these computations i.e. no ASVs were removed.

#### 5.5.1 Jensen Shannon divergence

Due to the compositional nature of the data, the composition of the samples can be viewed as probability distributions, summing to 1. The Jensen Shannon Divergence (JSD) (*94*) is symmetric, bounded and calculates distance between normalized probability dstributions. Here, it was calculated using Scipy’s (*79*) jensenshannon function. The JSD between two distributions *X* and *Y, J*_*X,Y*_ is defined in Equation 4.

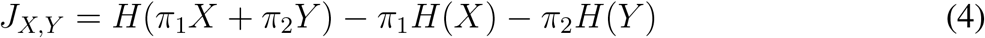

If *X* = {*x*_*i*_} and *Y* = {*y*_*i*_}, where *x*_*i*_ and *y*_*i*_ represent the normalized probability of finding the value *x*_*i*_ and *y*_*i*_ in the probability distributions *X* and *Y*, then the function *H* is the entropy of the distribution defined by Equation 5, similar to Equation 1.

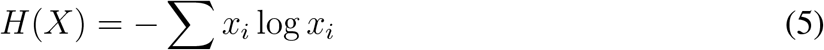

The weights *π*_1_ and *π*_2_ are set to be equal, i.e.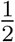. Using Equations 4 and 5, the Jensen Shannon divergence between the relative abundance composition of the two communities can be calculated. Note that the scipy function “jensenshannon” calculates the square root of Equation 4. These values are reported in Figure S9.

#### 5.5.2 Aitchison’s distance metric

Aitchison’s distance is a relevant metric for calculating distances between samples in compositional data (*89, 90*). A small pseudocount (0.001) was added to all the data to remove zeroes. The data were then transformed via clr. The Euclidean distance between the samples was then computed using the “euclidean” function from scipy. These distances are reported in Figure S11.

#### 5.5.3 Unifrac distance metric

To include the phylogenetic information in calculating the distances between communities, Unifrac distance metric was used. This metric needs a tree as the input. The same tree created for Faith’s PD measurement was used here. As before, the data set was reduced to contain only those taxa whose maximum relative abundance was atleast 0.5% in any sample. skbio’s unweighted unifrac function, which is based on (*86*), was used to compute the unifrac distances. These distances are reported in Figure S13.

#### 5.5.4 Embedding distances

In order to visualize distances between the communities better, Multi-dimensional embedding (MDS) was used to embed the distances in two dimensions. However, since a-priori the dimensionality is unknown, the stress of embedding was calculated was calculated for 1 to 5 dimensional spaces. These stresses are shown in the bottom panel of each embedding in Figures S10, S12 and S14. These show that the stress of a two dimensional embedding is close to the asymptotic value. Scikit-learn’s “MDS” function was used to perform the embedding. Note that each embedding was performed independently, so that coordinates across different embeddings cannot be compared.

### 5.6 Correlation networks

SCNIC (*43*) was used to obtain correlation modules and networks in the data. To enable interpretation, the data were thresholded to include only those taxa whose relative abundance was at least 5% in any community at any time point. This thresholded dataset was then subdivided into the three environments, paper, cellulose and glucose, and these were provided as inputs to the SCNIC package. As a first step, SCNIC uses a fast implementation of the package sparcc (*95*), to calculate correlations between all ASVs within each environment. sparcc takes into account the compoisitional nature of the data to estimate the correlation strengths. Next, SCNIC takes those ASVs whose correlation strength is 0.35 or higher, and groups them in modules so that each ASVs in the module is correlated to all other ASVs in the module via a correlation strength of 0.35 or higher. By such a definition, some ASVs cannot be placed in any modules and are indicated by light blue patches in the Files S1 and S2. Cytoscape (*96*) was used to create the network maps. Currently, SCNIC only works with positive correlations. In order to create modules of negative correlations, the signs of correlation strengths were inverted and this modified data was provided to SCNIC to create modules.

The correlation network data files were parsed using NetowrkX (*97*) in python to obtain the values of the strengths and number of correlations for comparing across environments.

### 5.7 Functional analysis

From the 16S sequence data, it is possible to get insights into the functional profiles of the communities. Here, two pipelines were used to obtain clues about the functions of the communities.

#### 5.7.1 PICRUSt2

The package PICRUSt2 (*44*) was used to infer Enzyme Classification (EC) numbers, Kegg Orthologs (KO) and pathway abundances in the samples. These will be referred to as ‘function types’. For all three cases, the unstratified output of PICRUSt2 was used, which provides abundances of each function per sample. In particular, this is a table for each function type where for each sample, the abundance of a function is estimated. Here, by function we mean E.C numbers for the function type Enzymes, KO numbers for the function type Kegg Orthologs and pathway names for the function type pathways.

##### Comparison across environments

For each function type, the abundance distribution of each function was compared between (1) all the cellulose broth communities and all the glucose broth communities and (2) all the cellulose paper communities and all the glucose broth communities. A two sample Kolmogorov-Smirnoff test was performed to estimate if the distribution of a function was significantly different between the two environments. For those functions that were significantly different, the difference in the median value of the function between the two environments was found. Based on this difference, the 15 functions that were high in cellulose broth and had the highest difference compared to glucose broth were chosen. Similarly, the 15 functions that were high in glucose and had the highest difference compared to cellulose broth were chosen. These are represented in Tables S1, S2, and S3. The functions selected from the comparison between cellulose paper and glucose were almost identical to those selected from the comparison between cellulose broth and glucose. Here only the latter are shown.

##### Distance metric

Euclidean distance was computed based on the abundance of functions between communities in a given environment, for each function type. Since the data are not compositional, the Euclidean metric can be directly applied. The entire dataset was used for this. The distance between replicates at each day is shown in Figures S25, S27, S29 for the function type enzymes, Kegg Orthologs and pathways respectively. The distances were embedded in 2 dimensions using the MDS method described above. These are shown in Figures S24, S26, S28 for the the function type enzymes, Kegg Orthologs and pathways respectively. Below each embedding, the stress of the MDS embedding is shown. The stress at 2 dimensions is similar to the stress at higher dimensions; so a 2 dimensional representation is acceptable in these cases.

##### Comparison of communities between the initial and the final days of the cycles in glucose broth

From Figures S24, S26, S28 it was noted that in the glucose broth, the communities in the initial days of all four Rounds clustered together (blue markers of all 4 types). To look for the differences in the function between the initial days and the final days, the communities at days 3 and 4 of all Rounds were grouped as the ‘initial days’ and the communities at days 13 and 14 of all Rounds were grouped as the ‘final days’. Days 3 and 4 were chosen instead of Days 1 and 2 to avoid transient dynamics. As before, for each function type, the abundance distribution of each function was compared between the initial days and the final days to check if they were significantly different between the two groups. The two sample Kolomogorov-Smirnoff test was used to test for significance, with a Bonferroni coreected p-value. For the functions that were significantly different, the median across the all communities in each group was found to know in which group the function is higher. Based on the difference in the medians, the functions were classified as being higher in the initial days and higher in the final days. This is shows in Table S1, S2 and S3 for the three function types.

##### Principal Component Analysis

PCA was performed on the data in the following way. First, the dataset was thresholded to remove very low abundance functions. The frequency of the maximum abundance of each function of each function type across all samples was plotted (Figure S21) and a threshold was set for each function type to remove those functions whose maximum abundance was very low across all samples. This reduced the number of functions: from 2413 to 610 for the enzymes, from 7802 to 1396 for the Kegg Orthologs, and from 438 to 361 for the pathways. Since these data are not compositional, first a PCA was performed directly on the dataset, and a distribution of the eigenvalues was found. Next, the data were randomized by shuffling the columns (functions) randomly for each row (samples). PCA was performed on this randomized dataset. However, on comparing the eigenvalue distribution, it was found that the highest eigenvalue remained in the randomized dataset, indicating higher order structure to the date. A careful analysis revealed that some of the samples had almost an order of magnitude higher total functions (e.g. total enzyme abundances) than others. To correct for this, the abundances were converted to relative abundances. PCA was then performed on the clr transformed data as described before. Now, when the eigenvalue distributions of the data was compared to the randomized data, the high eigenvalues disappeared (Figure S21C,F,I), indicating that the difference in total function abundances was indeed the reason why the highest eigenvalue was retained when the data were not compositional. Further, the eigenvalue distributions also showed that the first two principal components are the largest and could adequately describe the dataset. Figure 5A shows the PCA on the data with the function type enzymes. Figures S22 and S23 show the PCA on the data with function type Kegg Orthologs and pathways respectively. The percentage of variance explained is indicated on the axes of the plots.

Similar to the analysis of the 16S data, here the contribution of each function to the separation along PC1 was found for each function type. The top 98th percentile of functions that contribute to the separation were chosen. The clr of the abundances of these functions for all three function types in the cellulose broth and glucose broth environments are shown in Figure 5B, C and D.

#### 5.7.2 FAPROTAX

The package FAPROTAX (*45*) was used to analyze the 16s data. This package provides function predictions based on the 16S sequences. The output is a file that provides the abundance of each function in each community.

##### Comparison across environments

Similar to the analysis using PICRUSt2, the distribution of abundances of each function was compared between communities in (1) cellulose broth and glucose broth and (1) cellulose paper and glucose broth. The two sample Kolmogorov-Smirnoff test was used to test the null hypothesis that both samples were drawn from the same distribution. If the p-value was lower than the Bonferroni corrected threshold, then the null hypothesis was incorrect and the two samples are differently distributed. The medians across all samples within a group of the functions that were found to be differently distributed were found. Based on the difference in the medians, the functions were classified as either being higher in the cellulose broth environment or in the glucose environment. The functions are presented in Table S4. These were identical to the functions selected in the comparison between communities grown on cellulose paper and glucose broth.

##### Distance metric

Euclidean distances were computed between all pairs of communities as described above. The distance between the replicates for each day is shown in Figure S31. The distances within each environment were embedded on 2 dimensions using multi dimensional embedding, as shown in Figure S32. The stress of the embedding is indicated below each panel.

As can be seen from the embedding, for glucose the stress at 2 dimensions is considerable higher than at three dimensions. To check for clustering or separation at higher dimensions, the glucose communities were embedded in 3 dimensions. This is shown in Figure S33.

##### Comparison of communities between the initial and the final days of the transfer Rounds in glucose broth

As seen in Figure S32C, the glucose communities are clustered according to the day of growth across all transfer Rounds, i.e. there is a transition from blue to yellow markers. To investigate this further, the abundance distribution of each function was compared across two groups - the ‘initial days’ group consisting of all communities at Days 3 and 4, and the ‘final days’ group consisting of all communities at Days 13 and 14. As before, the first two days were not considered to avoid transient dynamics. The Kolmogorov-Smirnoff test was used, with a Bonferroni corrected p-value, to determine if the distribution of abundances was significantly different between the two groups. For those functions where it was different, the median of the function abundance was found across each group. The difference in medians between the two groups was used to determine in which group the function was dominant. This is shown in Table S4.

##### Principal Component Analysis

PCA was performed on the dataset, without any transformation of the data. The eigenvalue distribution is shown in Figure S30. The dataset was then randomized as describe above and PCA performed on the randomized dataaset. This showed that the two dominant eigenvalues disappeared on randomizing. So no further transformation of the data was needed. The results of the PCA are shown in Figure 6A-D.

In this case, the glucose and cellulose communities are separated along PC2. As described before, the contribution of each function to the separation along PC2 was found. The top 18 contributing functions were chosen. Their abundance across the glucose and cellulose communities are shown in Figure 6E.

## 6 Acknowledgements

The authors thank Dr. Ilya Pozdynakov, Prof. Alfred Spormann and Dr. Alan Derman for useful discussions on metabolism and John Sterret, University of Colorado, for help with SCNIC. This work was funded by the Max Planck Society.

## Author Contributions

K.H.P analyzed the data and wrote and revised the paper. Y.Z and P.B.R designed the study.

Y.Z. performed the experiments, acquired the data and revised the paper. A.D.F. and P.B.R. supervised the project and revised the paper.

---

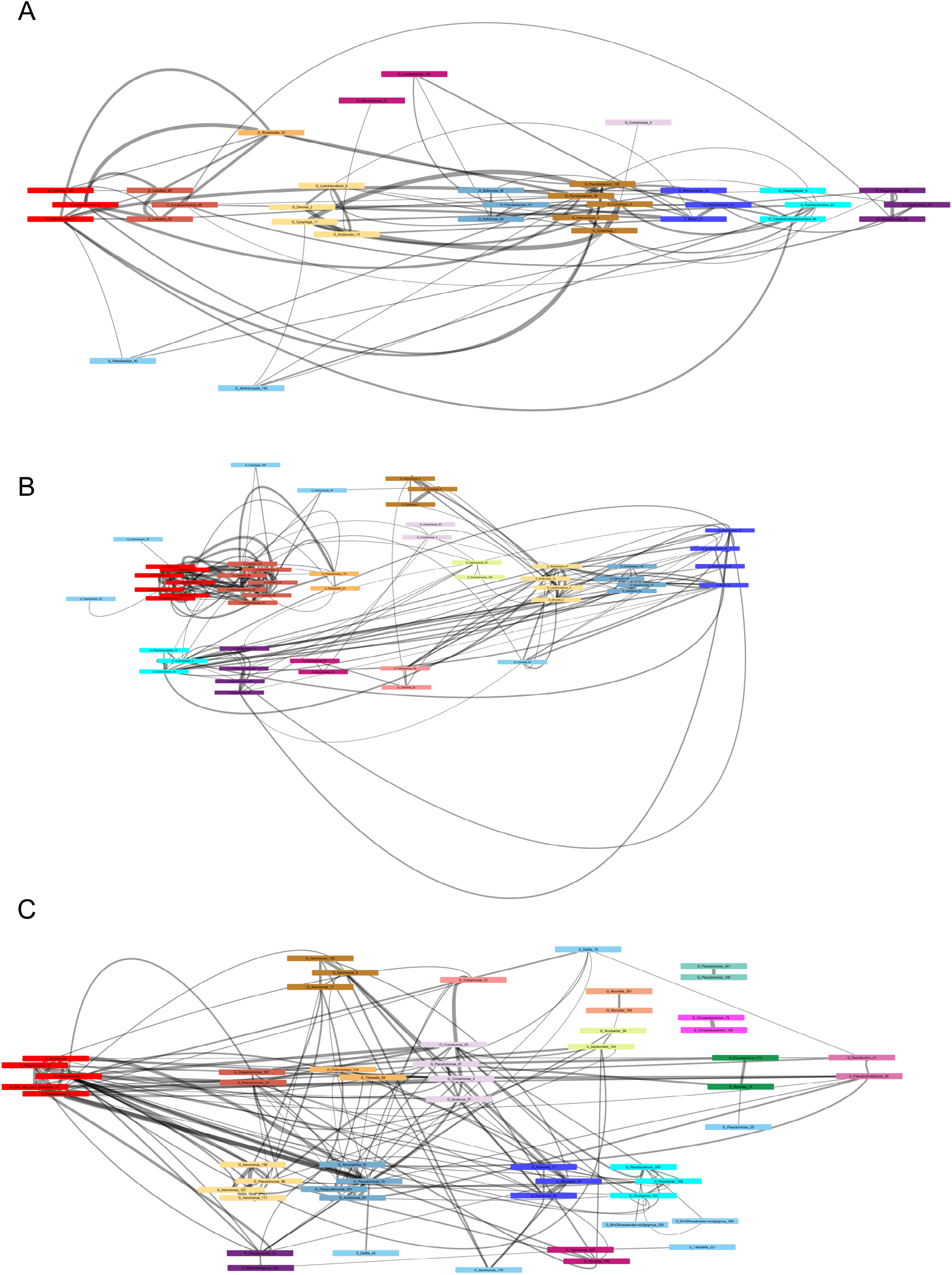

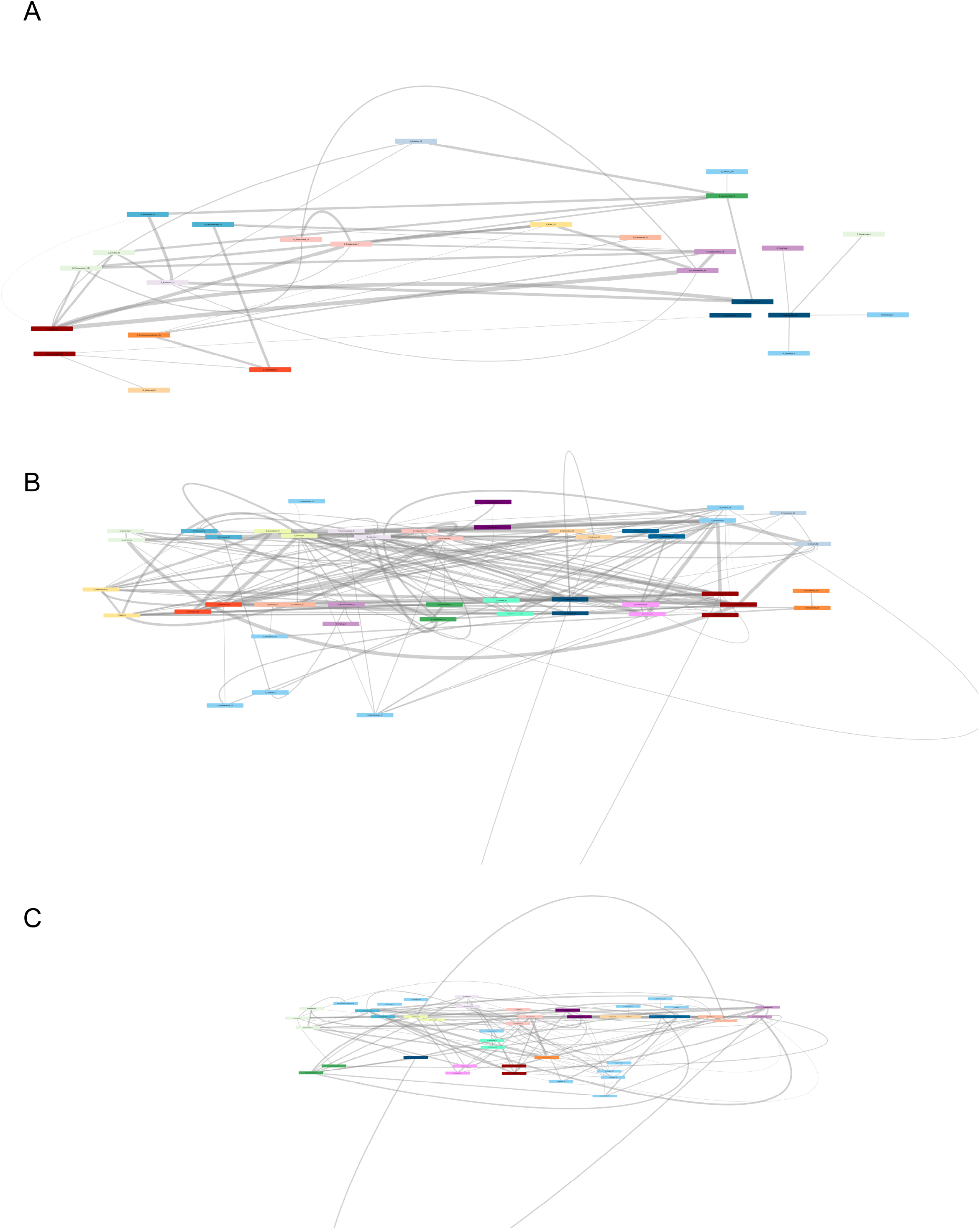

