## Supplementary_figures for "Diverse microbial communities assemble on both recalcitrant and labile carbon sources"

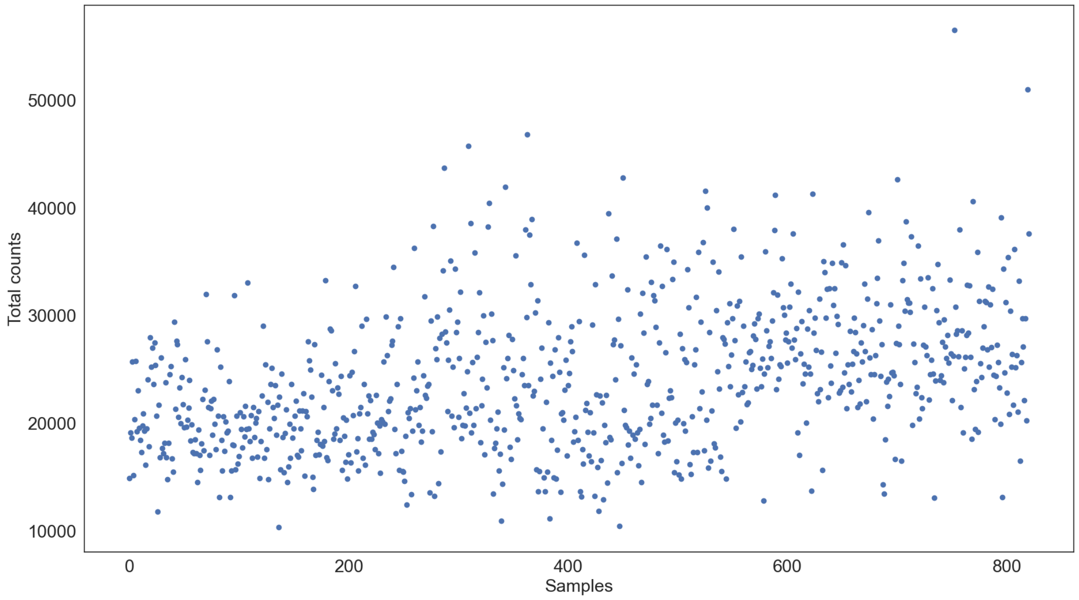


**Figure S1: Number of reads per sample:** The number of counts plotted on the y axis for each sample sequenced, plotted on the x axis.

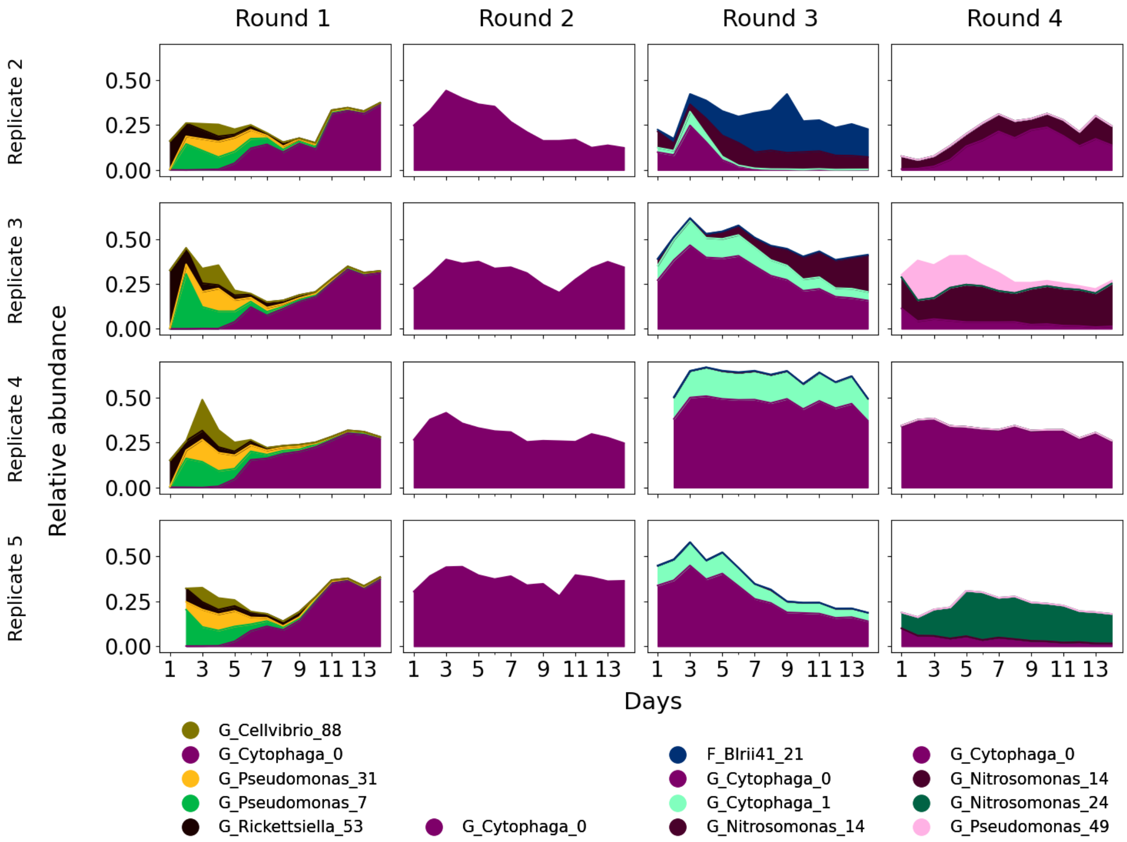


**Figure S2: Dynamics of dominant taxa in cellulose broth communities:** The dynamics of the dominant taxa in cellulose broth as defined in Methods for other 4 replicates are shown here. The y axis shows the relative abundance and the x axis shows the days of sampling. The columns are the rounds of transfer. Below each column the legends show the identity of the ASVs in terms of the lowest phylogenetic classification provided by SILVA. G denotes Genus, F denotes Family. The numbers at the end are unique identifiers for each ASV detected.

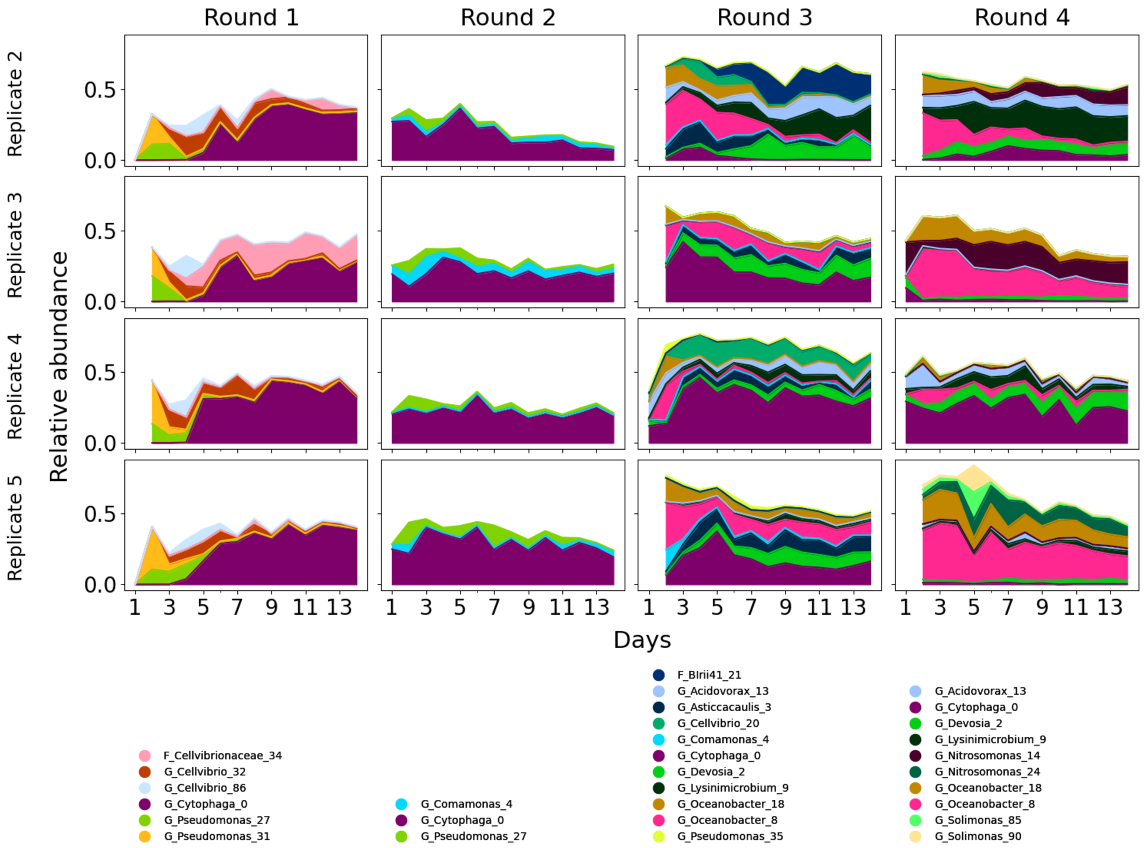


**Figure S3: Dynamics of dominant taxa on cellulose paper communities:**.The dynamics of the dominant taxa on cellulose paper as defined in Methods for other 4 replicates are shown here. The y axis shows the relative abundance and the x axis shows the days of sampling. The columns are the rounds of transfer. Below each column the legends show the identity of the ASVs in terms of the lowest phylogenetic classification provided by SILVA. G denotes Genus, F denotes Family. The numbers at the end are unique identifiers for each ASV detected.

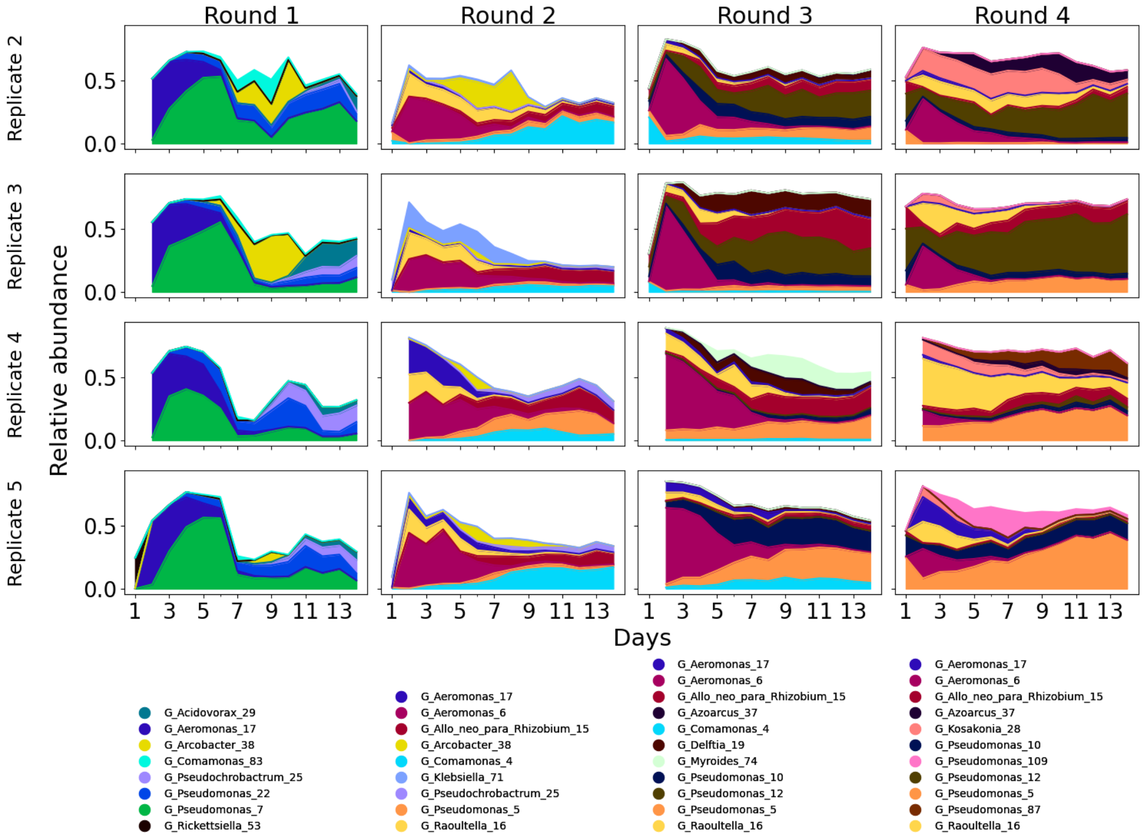


**Figure S4: Dynamics of dominant taxa in glucose communities:** The dynamics of the dominant taxa in glucose as defined in Methods for other 4 replicates are shown here. The y axis shows the relative abundance and the x axis shows the days of sampling. The columns are the rounds of transfer. Below each column the legends show the identity of the ASVs in terms of the lowest phylogenetic classification provided by SILVA. G denotes Genus. The numbers at the end are unique identifiers for each ASV detected.

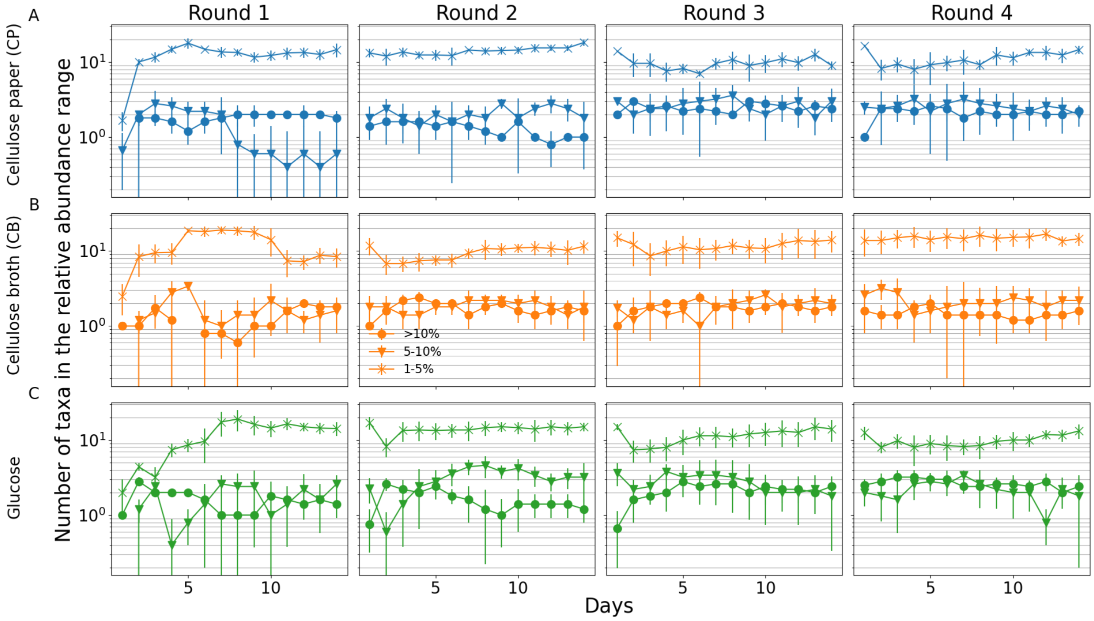


**Figure S5: Number of taxa within given relative abundance ranges:** The number of taxa in relative abundance ranges above 10%, denoted by circles, between $5-10\%$ denoted by triangles, and between $1-5\%$ denoted by asterisks, are plotted by averaging the abundances across replicates. The top row shows these data for communities in cellulose broth, the middle row for communities on paper, and the bottom row for communities on glucose.

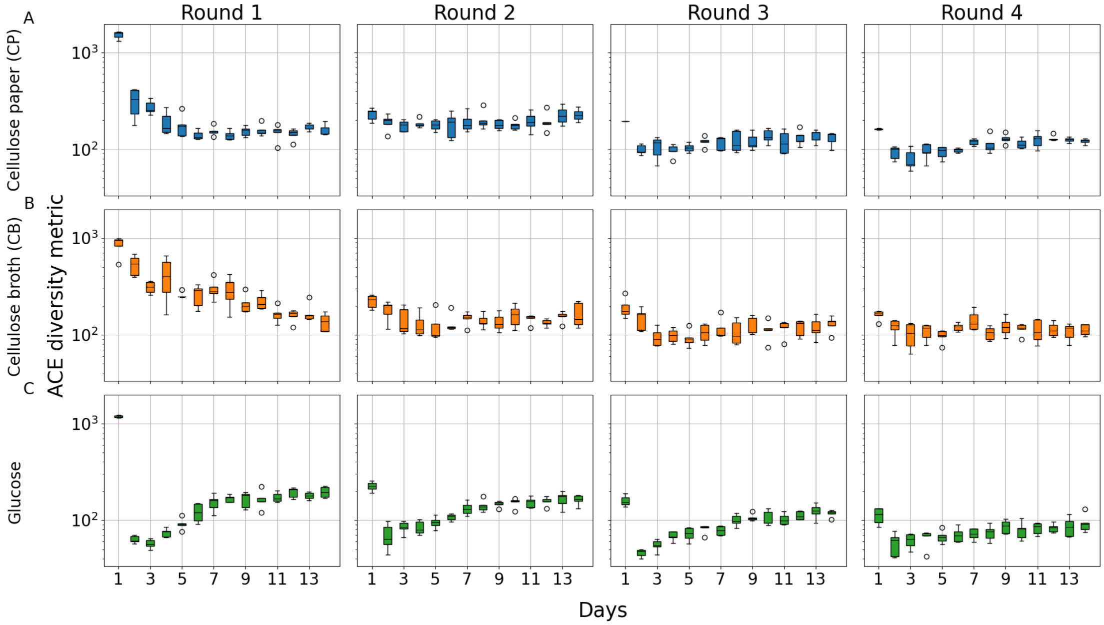


**Figure S6: ACE diversity metric:** The alpha diversity metric, Abundance based Coverage Estimate, which represents the richness, is shown on the y axis. The x axis shows the days of sampling. Each column shows the rounds of transfer. The top row shows these data for communities in cellulose broth, the middle row for communities on paper, and the bottom row for communities on glucose.

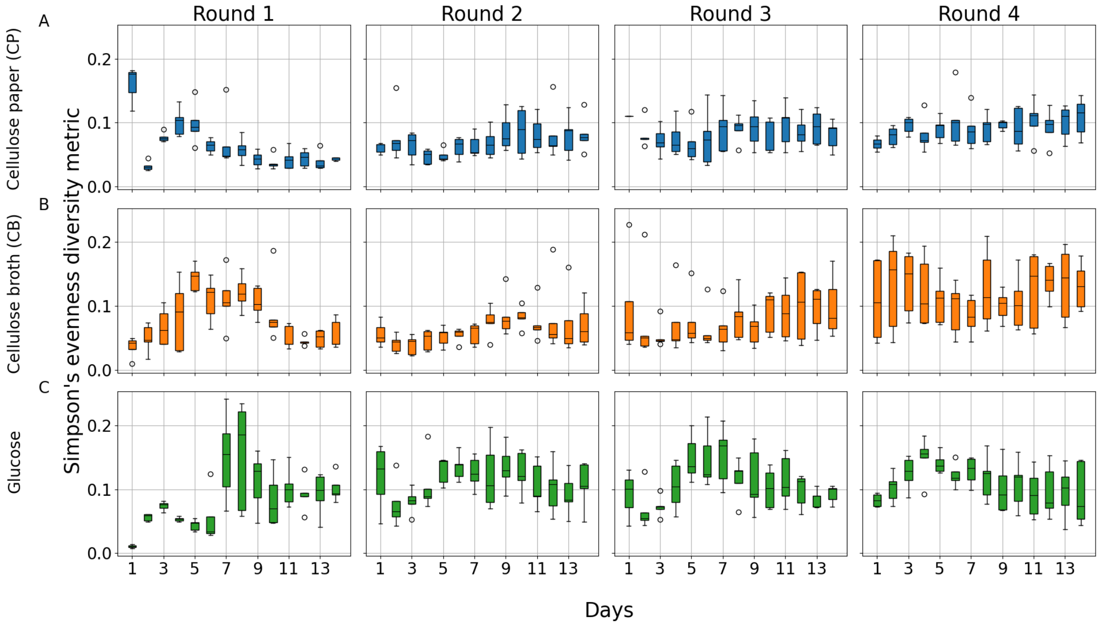


**Figure S7: Simpson's evenness measure:** The alpha diversity metric, Simpson's evenness, which represents the evenness, is shown on the y axis. The x axis shows the days of sampling. Each column shows the rounds of transfer. The top row shows these data for communities in cellulose broth, the middle row for communities on paper, and the bottom row for communities on glucose.

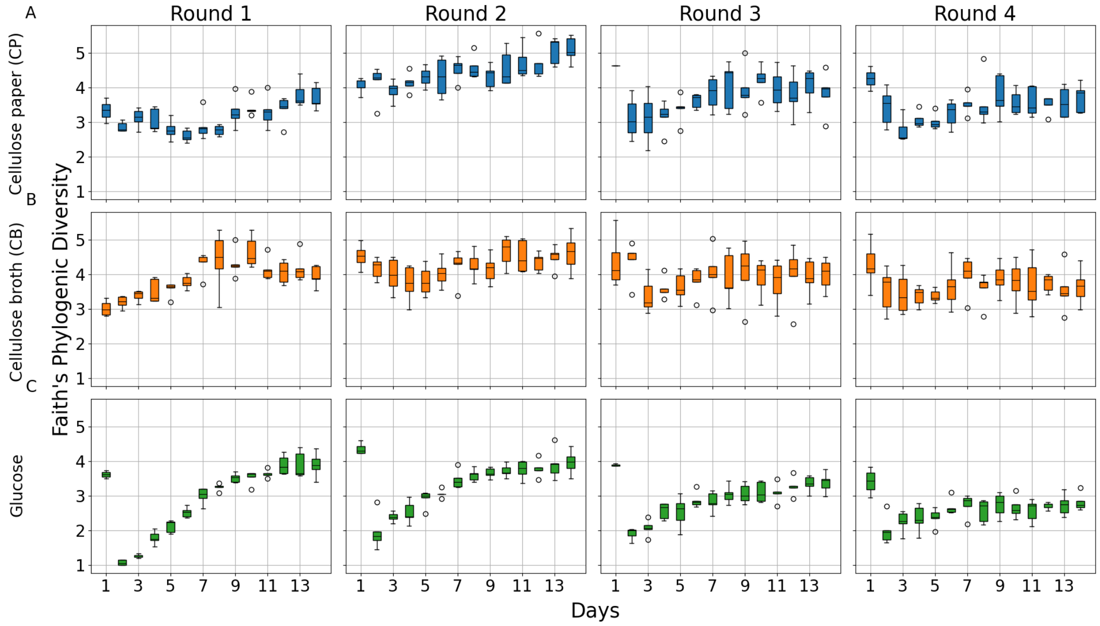


**Figure S8: Faith's phylogenetic diversity:** The alpha diversity metric, Faith's PD, which takes into account the phylogenetic information, is shown on the y axis. The x axis shows the days of sampling. Each column shows the rounds of transfer. The top row shows these data for communities in cellulose broth, the middle row for communities on paper, and the bottom row for communities on glucose.

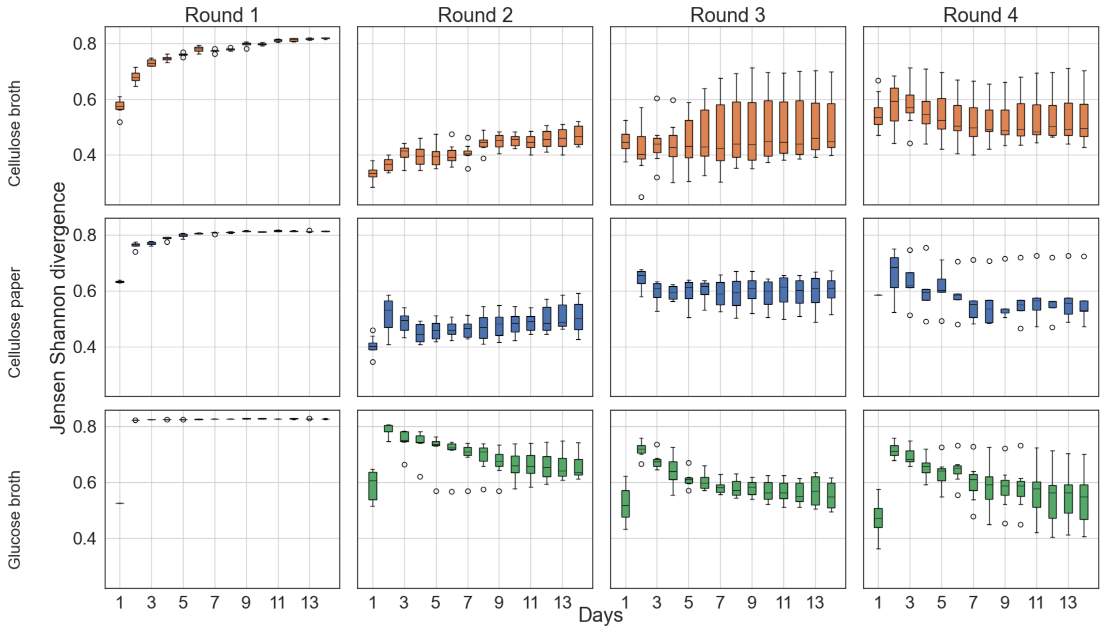


**Figure S9: Jensen Shannon divergence (JSD):** The beta diversity metric, JSD, is plotted on the y axis for the replicates. The x axis shows the days of sampling. Each column shows the rounds of transfer. The top row shows these data for communities in cellulose broth, the middle row for communities on paper, and the bottom row for communities on glucose. Each boxplot represents values across 5 replicate communities in each environment, where the boxes extend from the first to the third quartile, solid line represents the median, and the whiskers extend from the box to the farthest data point lying within 1.5 times the inter-quartile range.

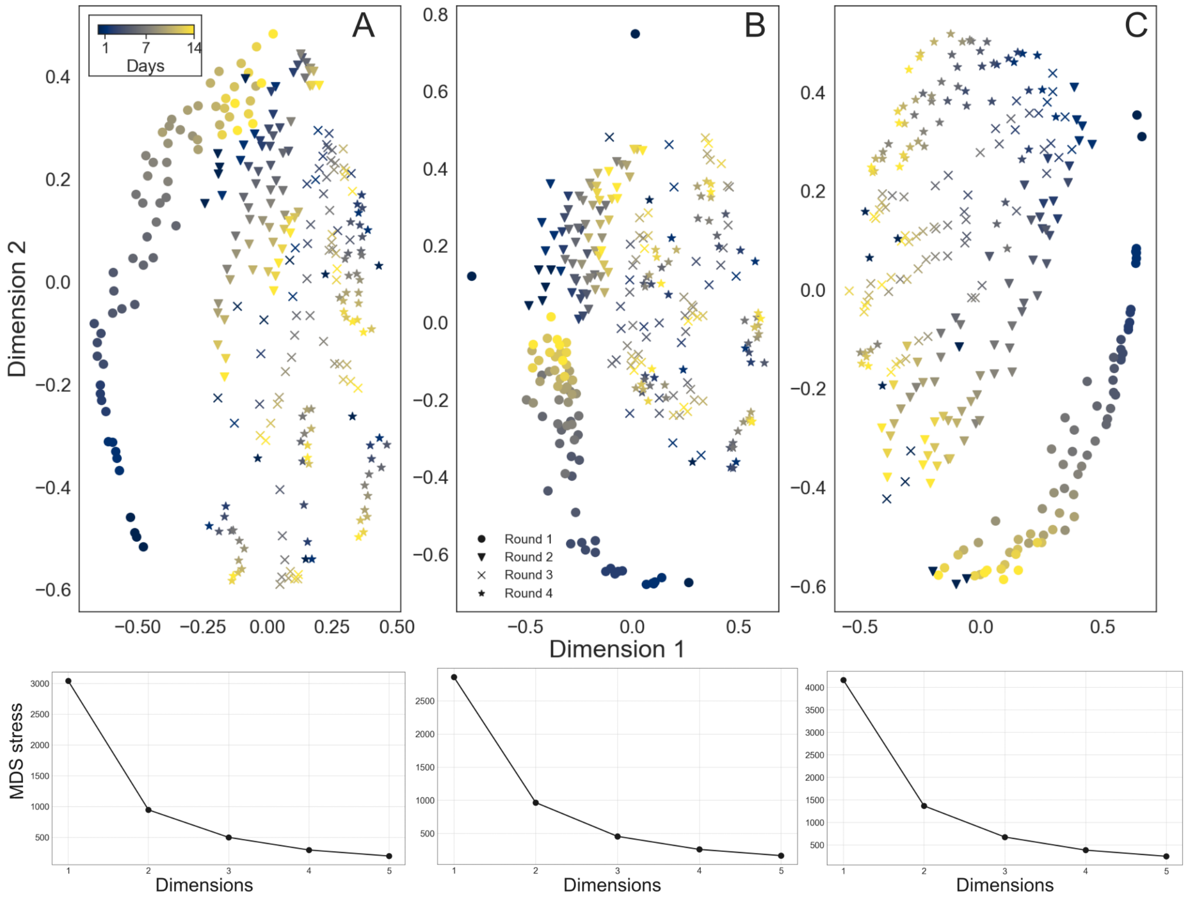


**Figure S10: Embedding of Jensen Shannon Divergence metric:** The JSD between all samples were embedded in two dimensions for each environment. **A** for cellulose broth **B** for paper and **C** for glucose. The bottom panels show the stress of the MDS embedding in each environment.

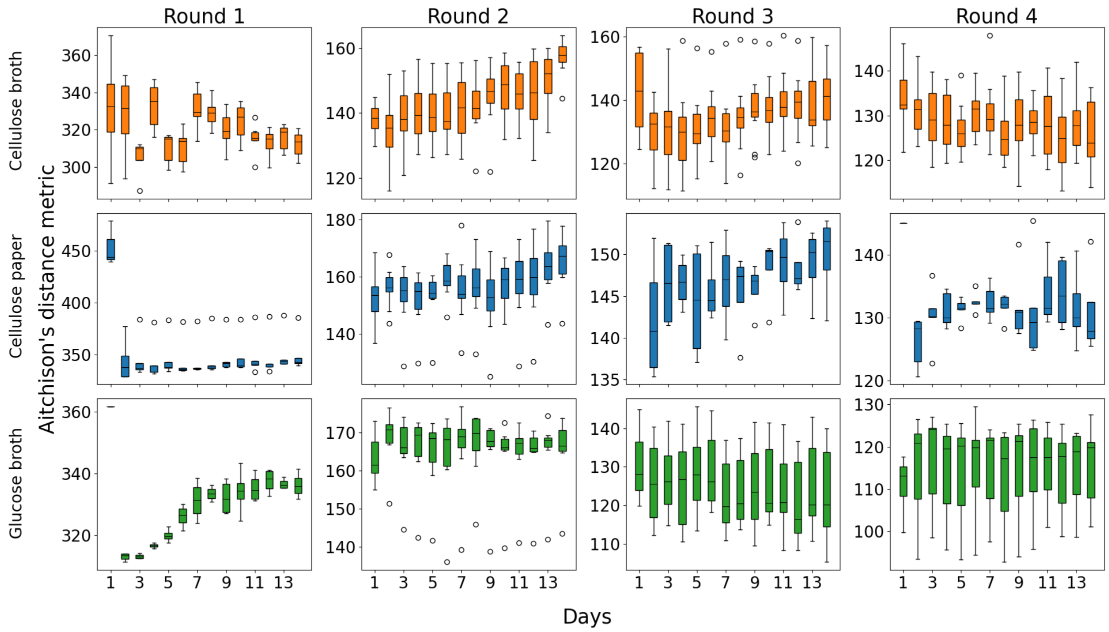


**Figure S11: Aitchison's distance metric:** The beta diversity metric, Aitchison's metric, which takes into account the compositional nature of the data is plotted on the y axis for the replicates. The x axis shows the days of sampling. Each column shows the rounds of transfer. The top row shows these data for communities in cellulose broth, the middle row for communities on paper, and the bottom row for communities on glucose. Each boxplot represents values across 5 replicate communities in each environment, where the boxes extend from the first to the third quartile, solid line represents the median, and the whiskers extend from the box to the farthest data point lying within 1.5 times the inter-quartile range.

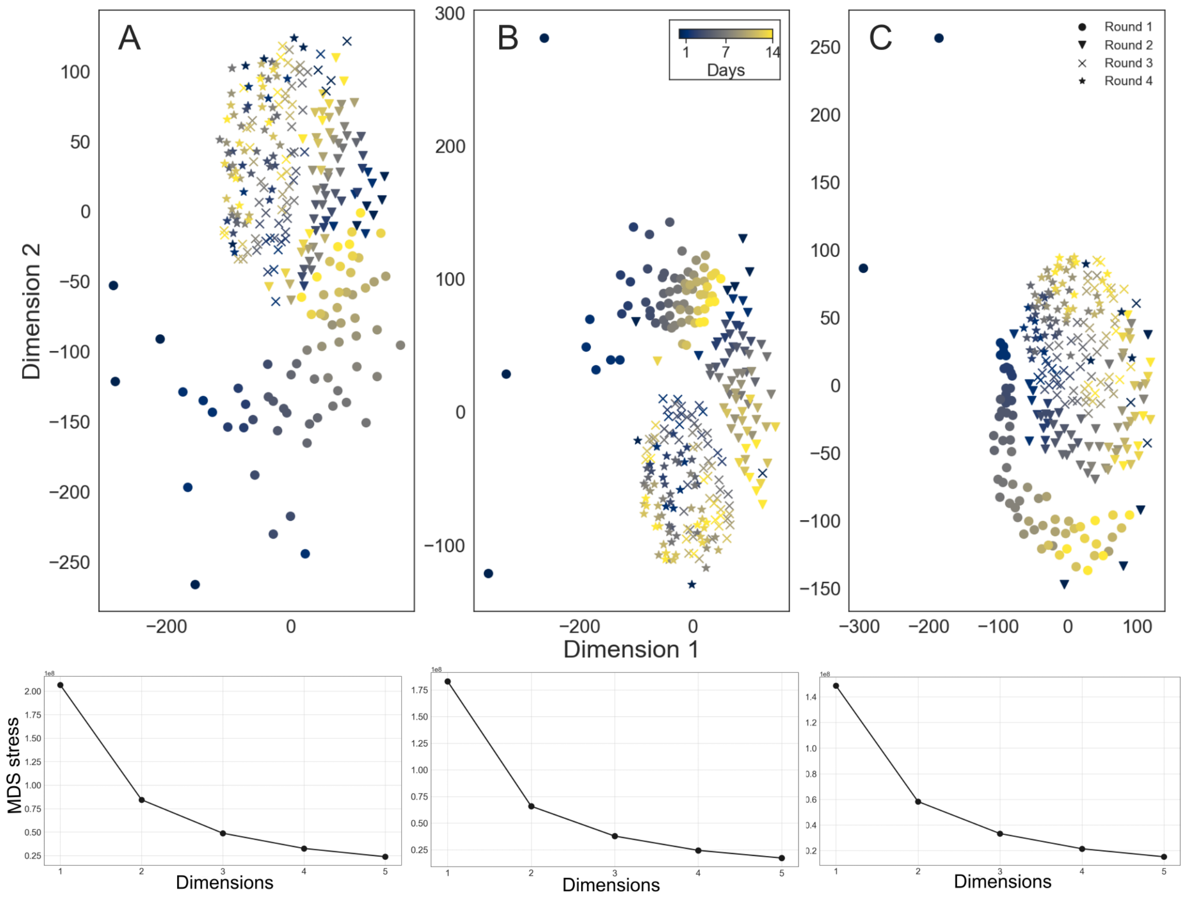


**Figure S12: Embedding Aitchison's distances:** The Aitchison's distance between all samples were embedded in two dimensions for each environment. **A** for cellulose broth **B** for paper and **C** for glucose. The bottom panels show the stress of the MDS embedding in each environment.

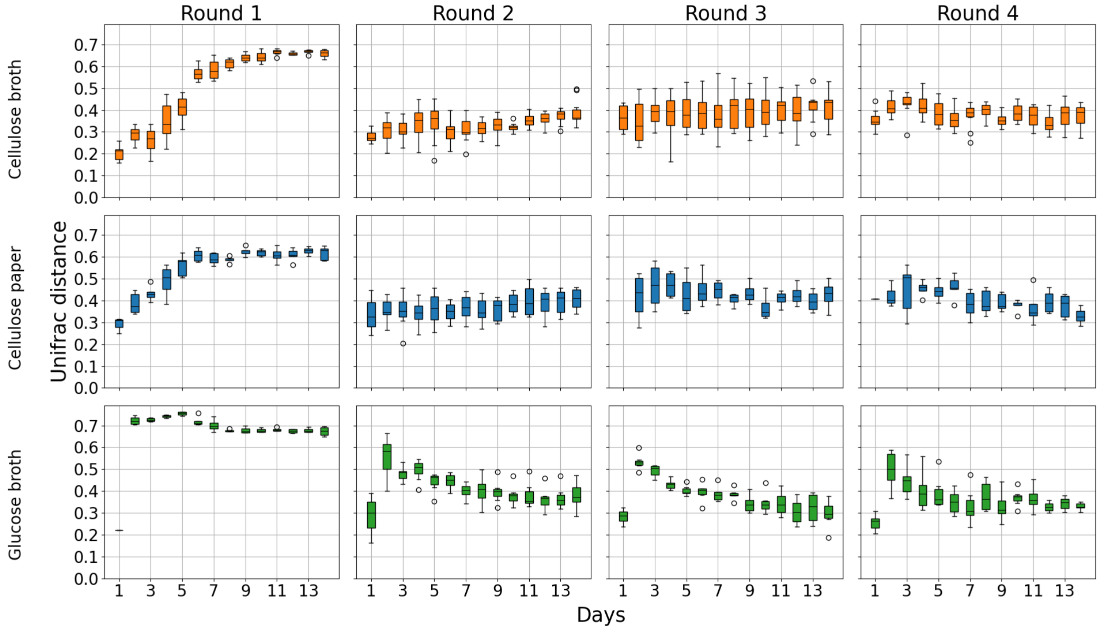


**Figure S13: Unifrac distances across days:** The beta diversity metric, Unifrac, which takes into account the phylogenetic information of the samples is plotted on the y axis for the replicates. The x axis shows the days of sampling. Each column shows the rounds of transfer. The top row shows these data for communities in cellulose broth, the middle row for communities on paper, and the bottom row for communities on glucose. Each boxplot represents values across 5 replicate communities in each environment, where the boxes extend from the first to the third quartile, solid line represents the median, and the whiskers extend from the box to the farthest data point lying within 1.5 times the inter-quartile range.

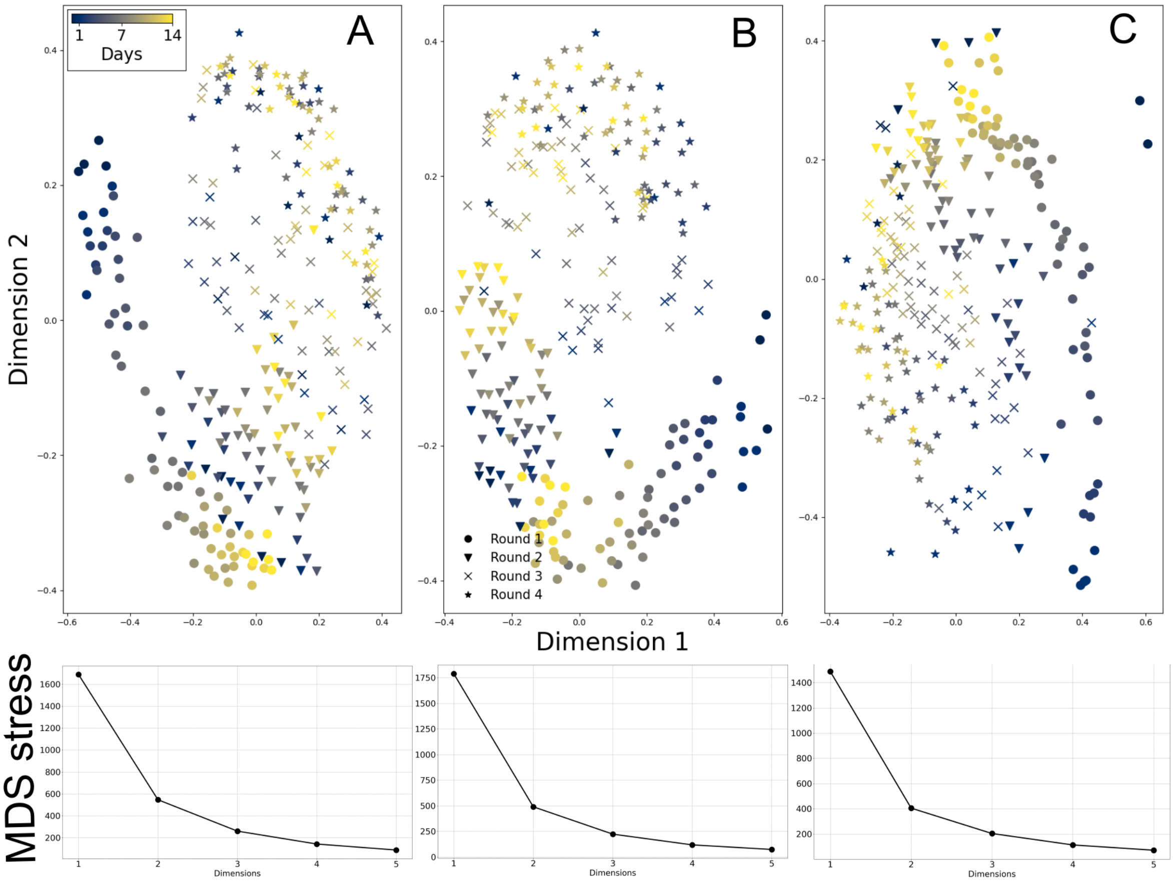


**Figure S14: Embedding Unifrac distances:**.The Unifrac distance between all samples were embedded in two dimensions for each environment. **A** for cellulose broth **B** for paper and **C** for glucose. The bottom panels show the stress of the MDS embedding in each environment.

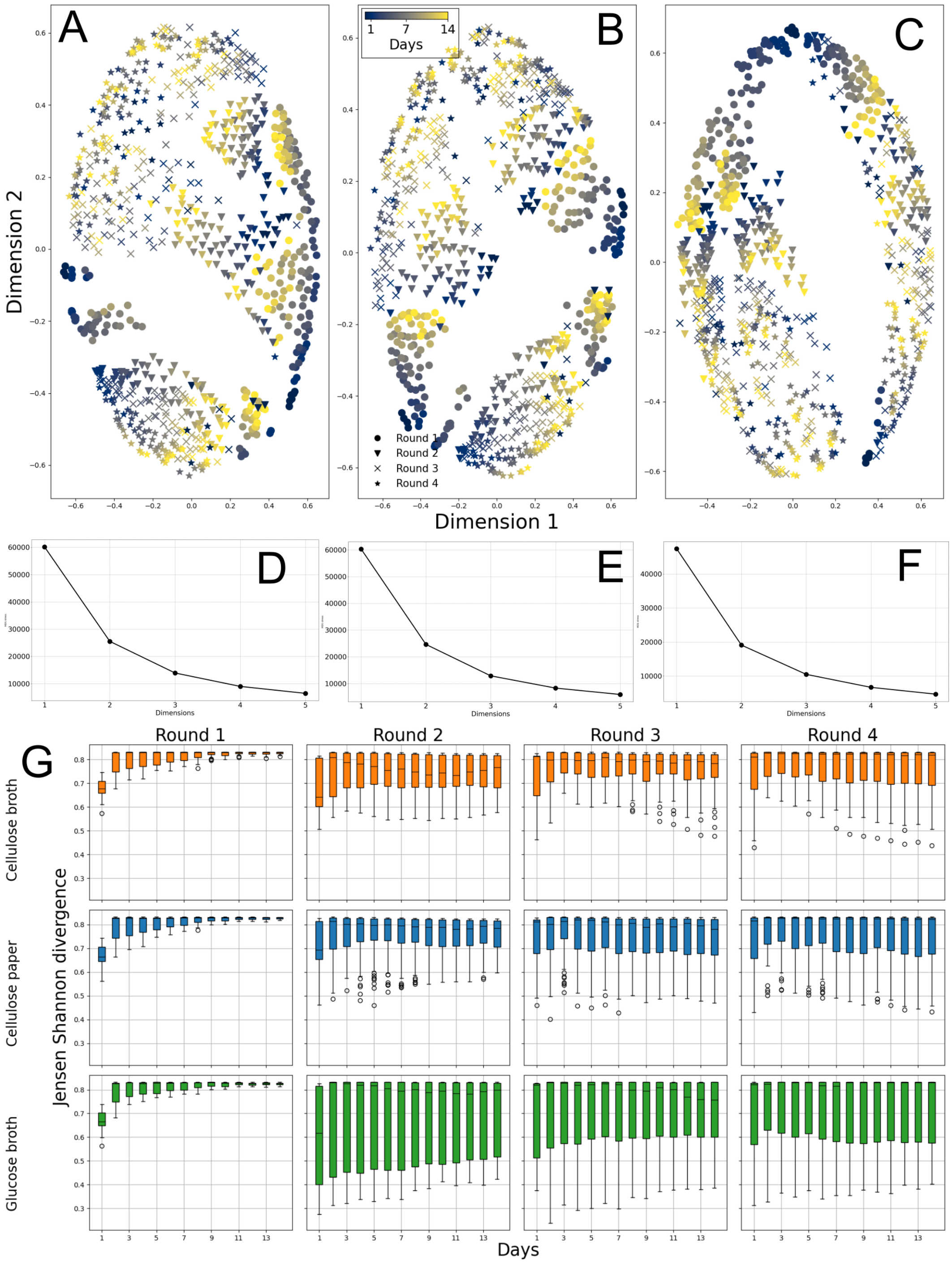


**Figure S15: JSD for only low abundance taxa:** The JSD was computed for the low abundance ASVs as described in methods. The pairwise distances were embedded in two dimensions for each environment. **A** for cellulose broth **B** for paper and **C** for glucose. **D-F** show the stress of the MDS embedding in each environment. **G** shows the distances between replicates; the top row for cellulose broth, the middle row for cellulose paper and the bottom row for glucose broth. Each boxplot represents values across 5 replicate communities in each environment, where the boxes extend from the first to the third quartile, solid line represents the median, and the whiskers extend from the box to the farthest data point lying within 1.5 times the inter-quartile range.

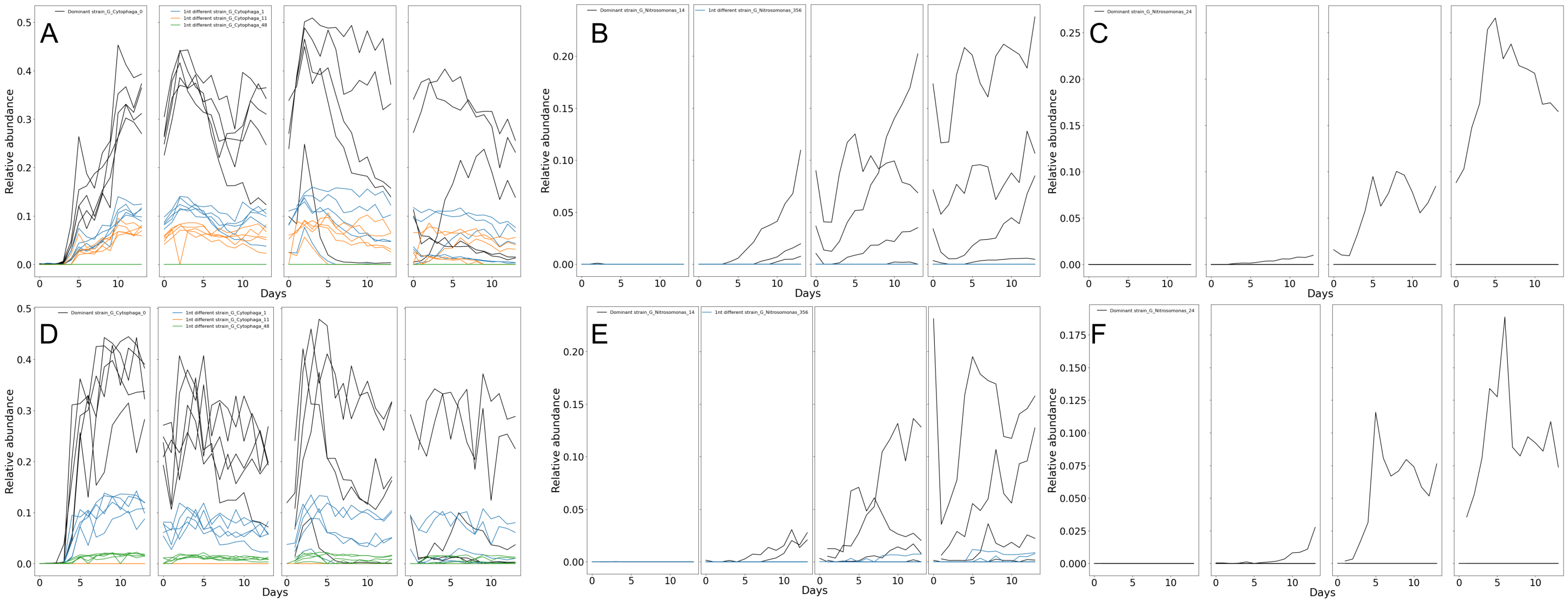


**Figure S16: ASVs with 1 nucleotide differences in the cellulose environment:** The relative abundance of the dominant ASV is plotted in black across all 14 days of all 4 rounds. In other colors, relative abundance of ASVs that are 1 nucleotide different from the dominant ASV are plotted **A-C** in the cellulose broth, **D-F** on paper. The different curves represent abundances in different replicates.

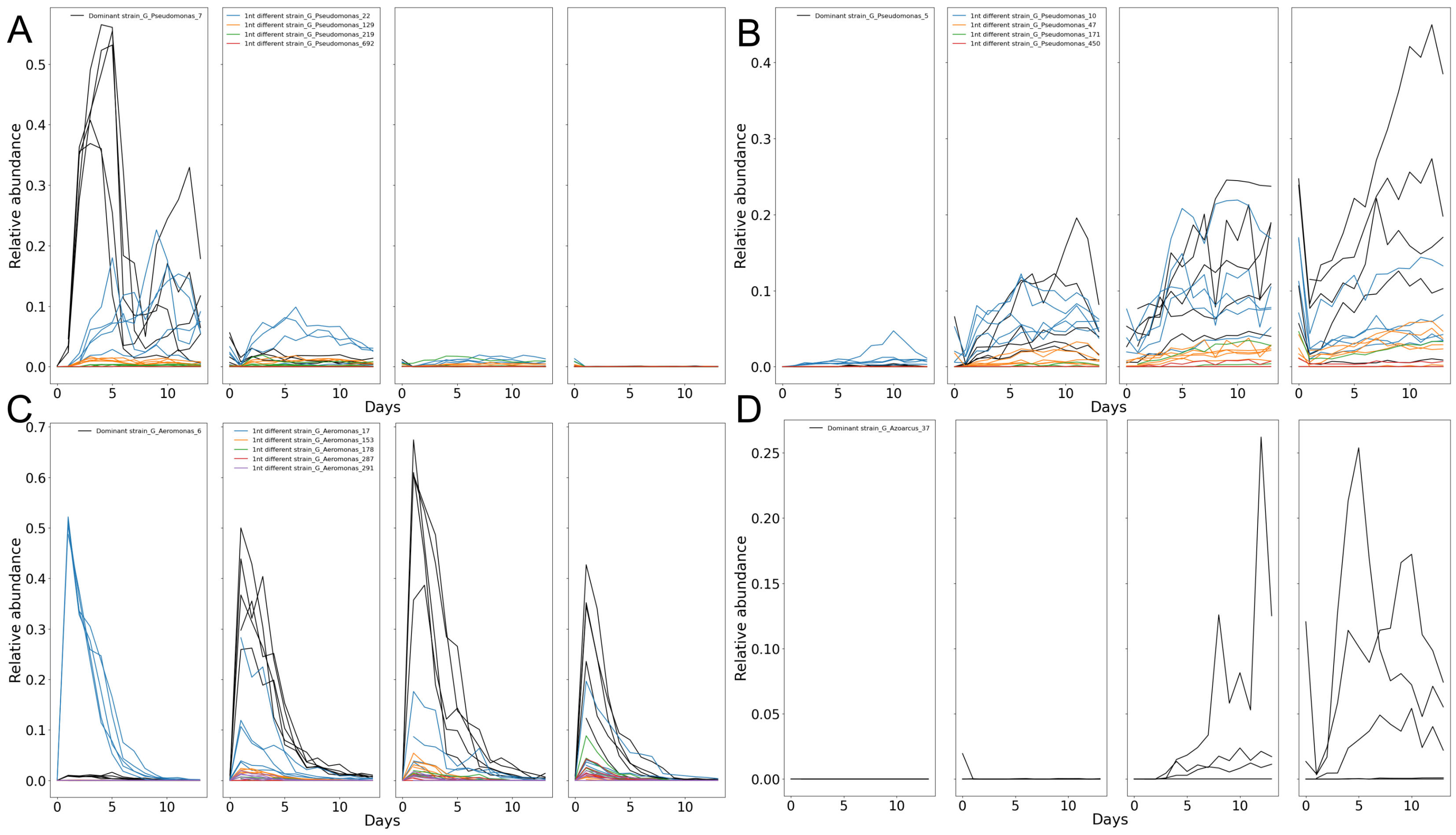


**Figure S17: ASVs with 1 nucleotide differences in the glucose environment:**The relative abundance of the dominant ASV is plotted in black across all 14 days of all 4 rounds. In other colors, relative abundance of ASVs that are 1 nucleotide different from the dominant ASV are plotted for **A** Psedumonas 7, **B** Pseudmonas 5, **C** Aeromonas 6 and **D** Azoarcus 37. The different curves represent the abundances in different replicates.

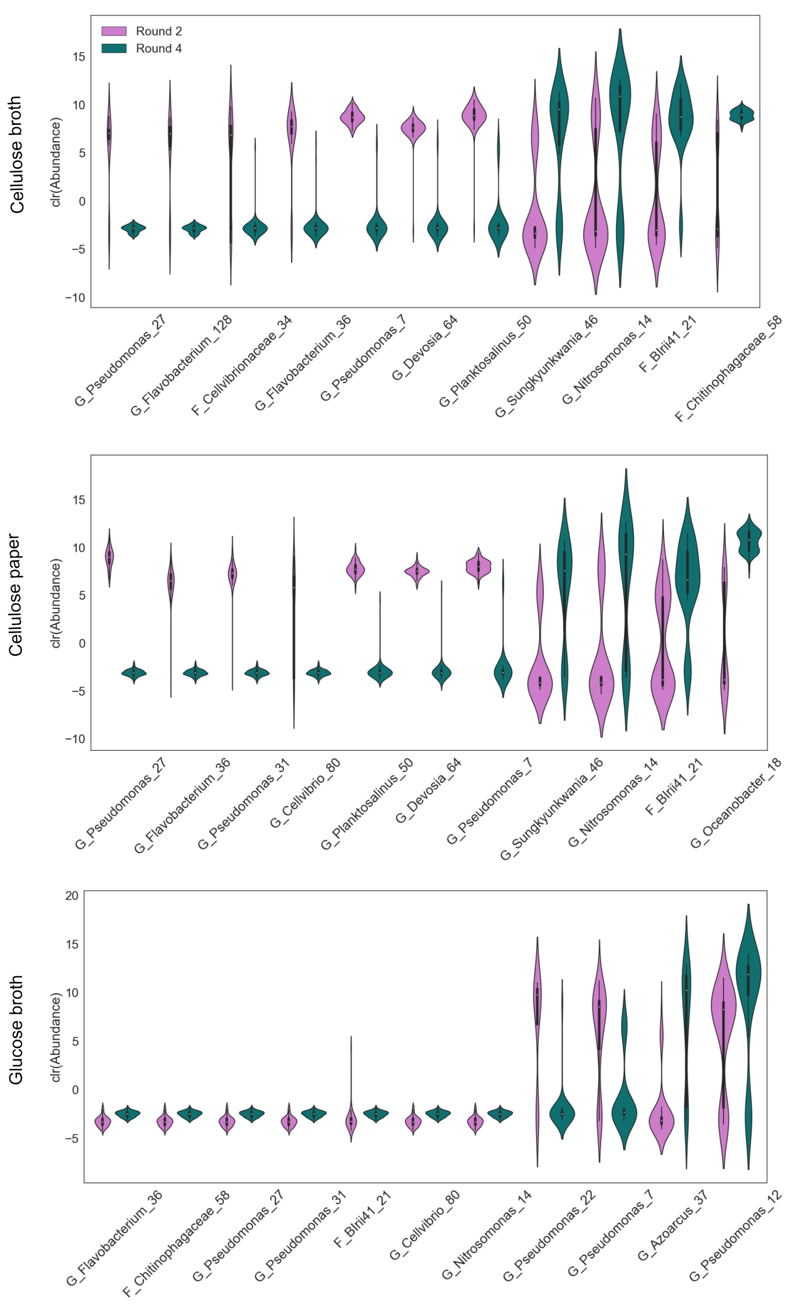


**Figure S18: ASVs causing displacement of communities along PC2 between Round 2 and Round 4:** The contribution of the ASVs to the displacement of communities between rounds 2 and 4 were calculated the top contributors were chosen, as described in Methods. The log transformed relative abundances in rounds 2 and 4 for these top contributors are plotted for the cellulose broth environment in the top panel, the cellulose paper environment for the middle panel and the glucose broth environment for the bottom panel. The violin plots represent the distributions across the 5 technical replicates.

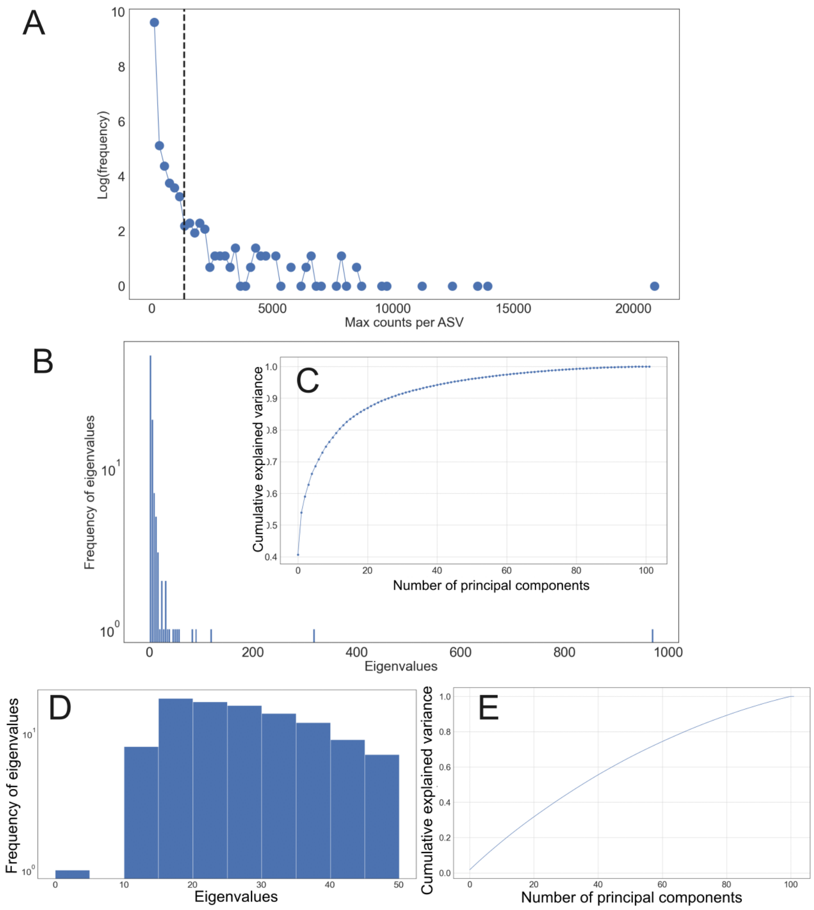


**Figure S19: Thresholding and eigenvalues of thresholded data:** For the PCA on 16S sequence data, the thresholding was performed as in Methods. **A** shows the logarithm of the frequency of maximum counts an ASV has across all sample. The dashed line shows the cutoff for thresholding. **B** The eigenvalue distribution for the thresholded data. **C** the cumulative explained variance **D** The eigenvalue distribution and **E** cumulative explained variance of the shuffled data.

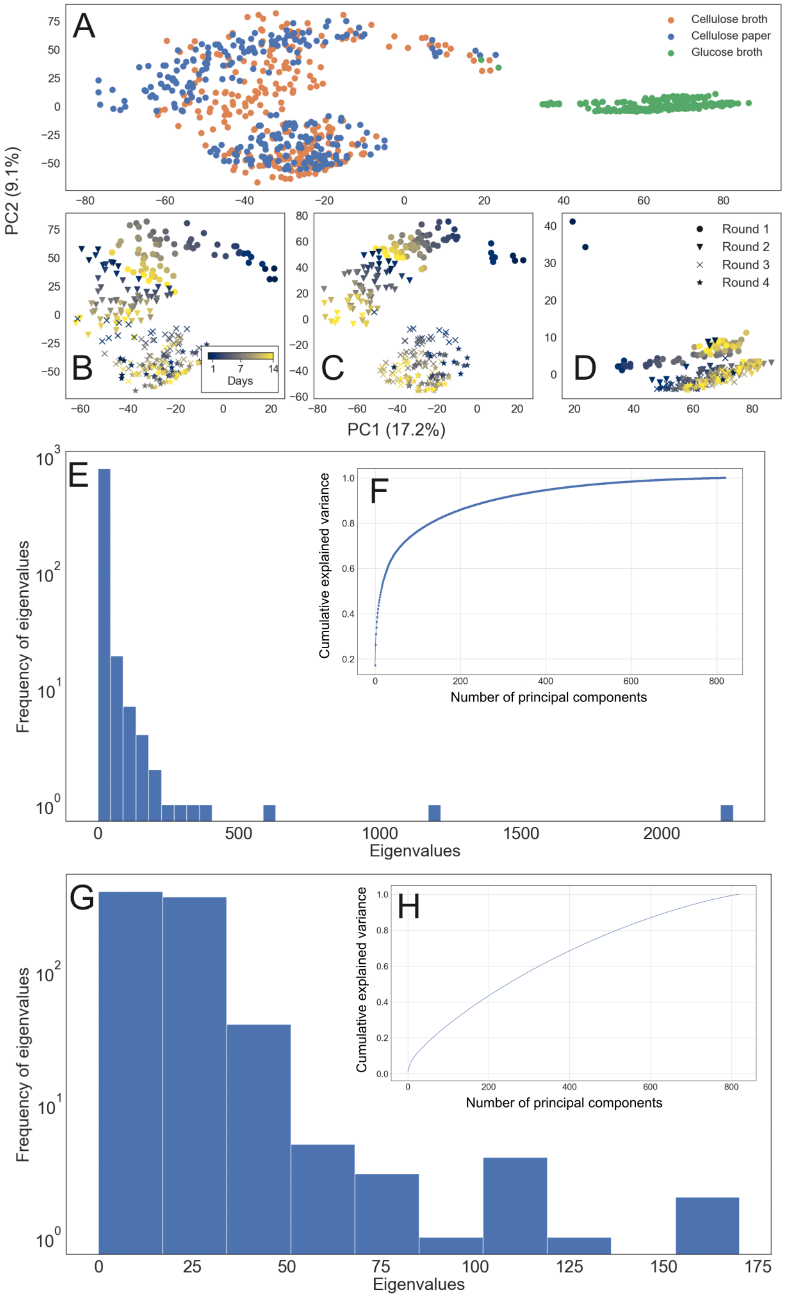


**Figure S20: PCA with the full data set: A** PCA embedded in two dimensions on the full 16S data. The colors represent the environment. **B-D** The same embedding as in A, but separated by the environments: cellulose broth, paper and glucose respectively. The colors represent the 14 days and the markers represent the 4 rounds of transfer. **E** Eigenvalue distribution **F** Cumulative explained variance **G** Eigenvalue distribution and **H** cumulative explained variance of shuffled data.

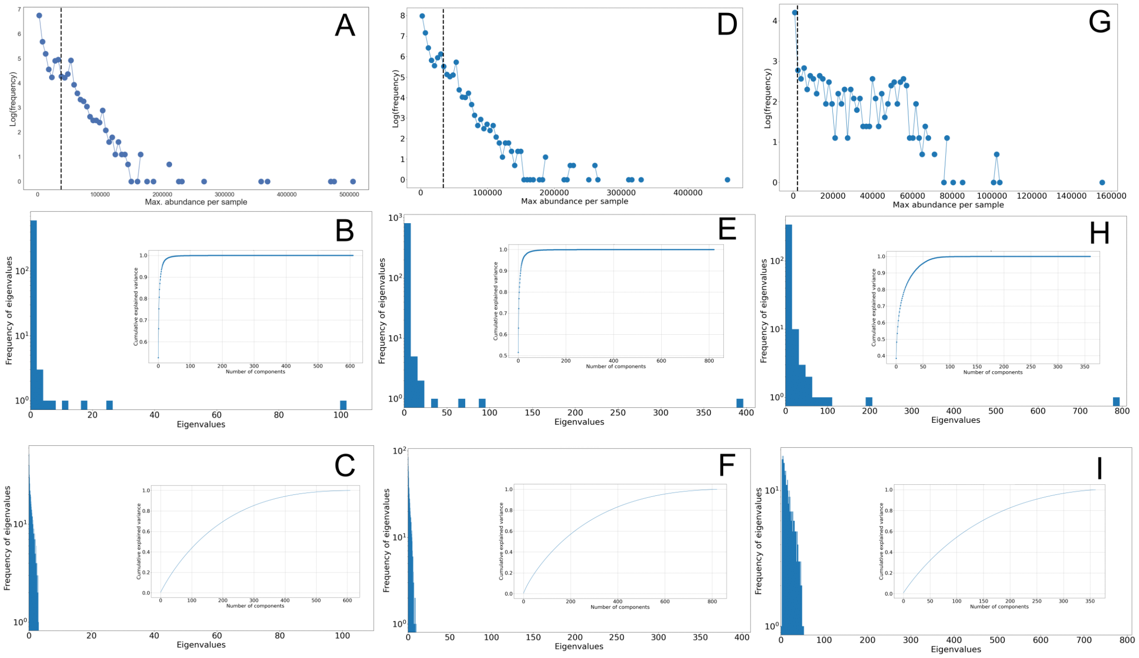


**Figure S21: Details of PCA on PICRUSt2 output:** PICRUSt2 provides 3 outputs - abundances of EC number, KO and pathways. The logarithm of the frequency of the maximum abundance across all samples, the eigenvalue distribution with an inset of the cumulative explained variance, and the eigenvalue distribution and cumulative explained variance of the shuffled data are shown for **A-C** cellulose broth environment **D-F** cellulose paper and **G-I** glucose.

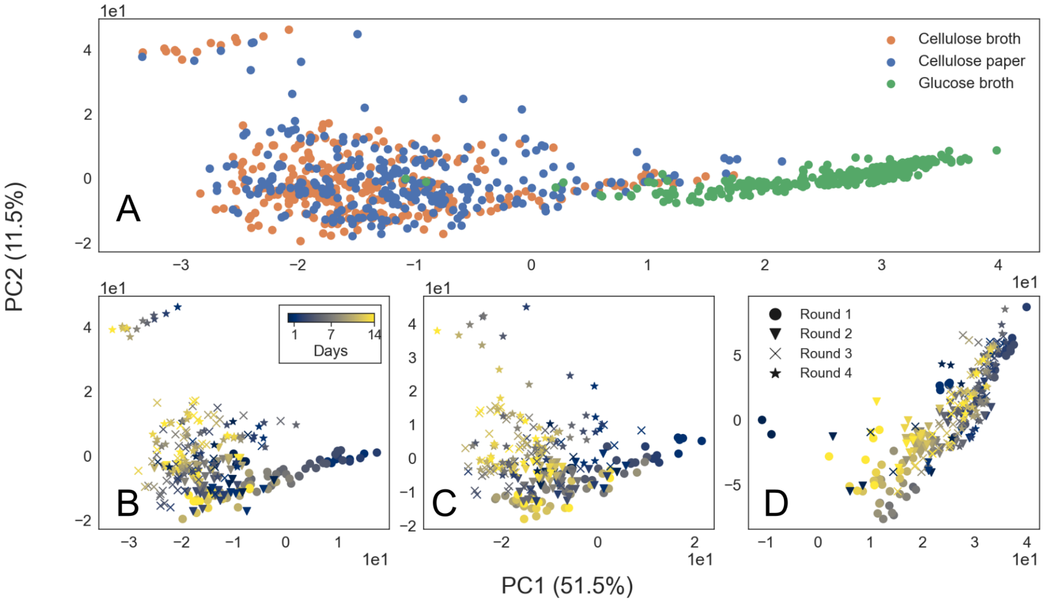


**Figure S22: PCA on Kegg Ortholog abundance output of PICRUSt2:** PCA was performed on the KO abundances as described in methods. **A** shows the 2 dimensional embedding of the sample, with the colors corresponding tot he environment. **B-D** shows the same embedding as in A, but separated by the environments: cellulose broth, paper and glucose respectively. The colors show the 14 days of sampling within each round, and the markers represent the rounds of transfer.

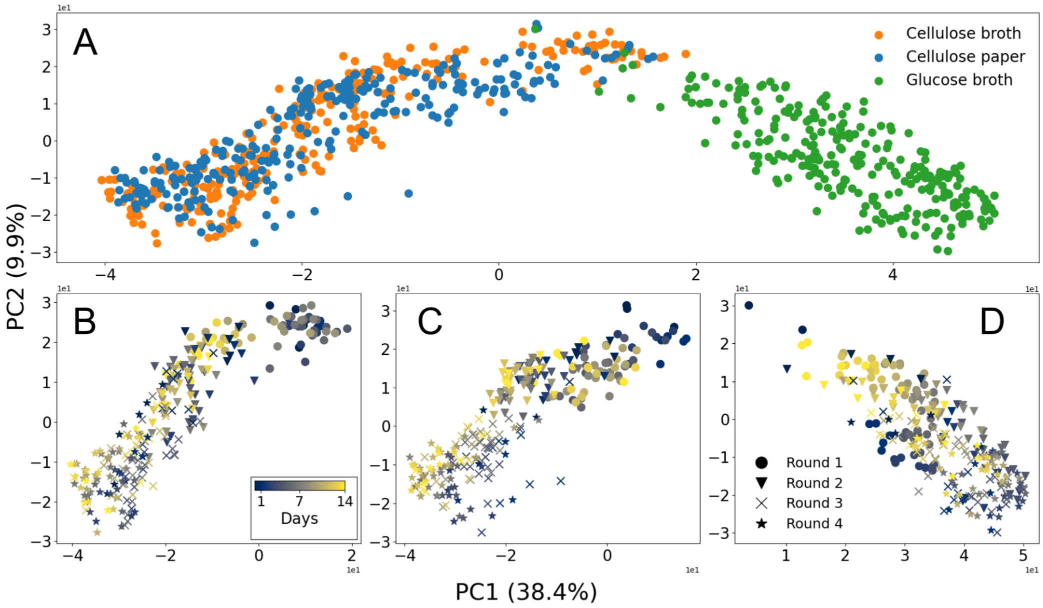


**Figure S23: PCA on pathway abundance output of PICRUSt2:** PCA was performed on the pathway abundances as described in methods. **A** shows the 2 dimensional embedding of the sample, with the colors corresponding to the environment. **B-D** shows the same embedding as in A, but separated by the environments: cellulose broth, paper and glucose respectively. The colors show the 14 days of sampling within each round, and the markers represent the rounds of transfer.

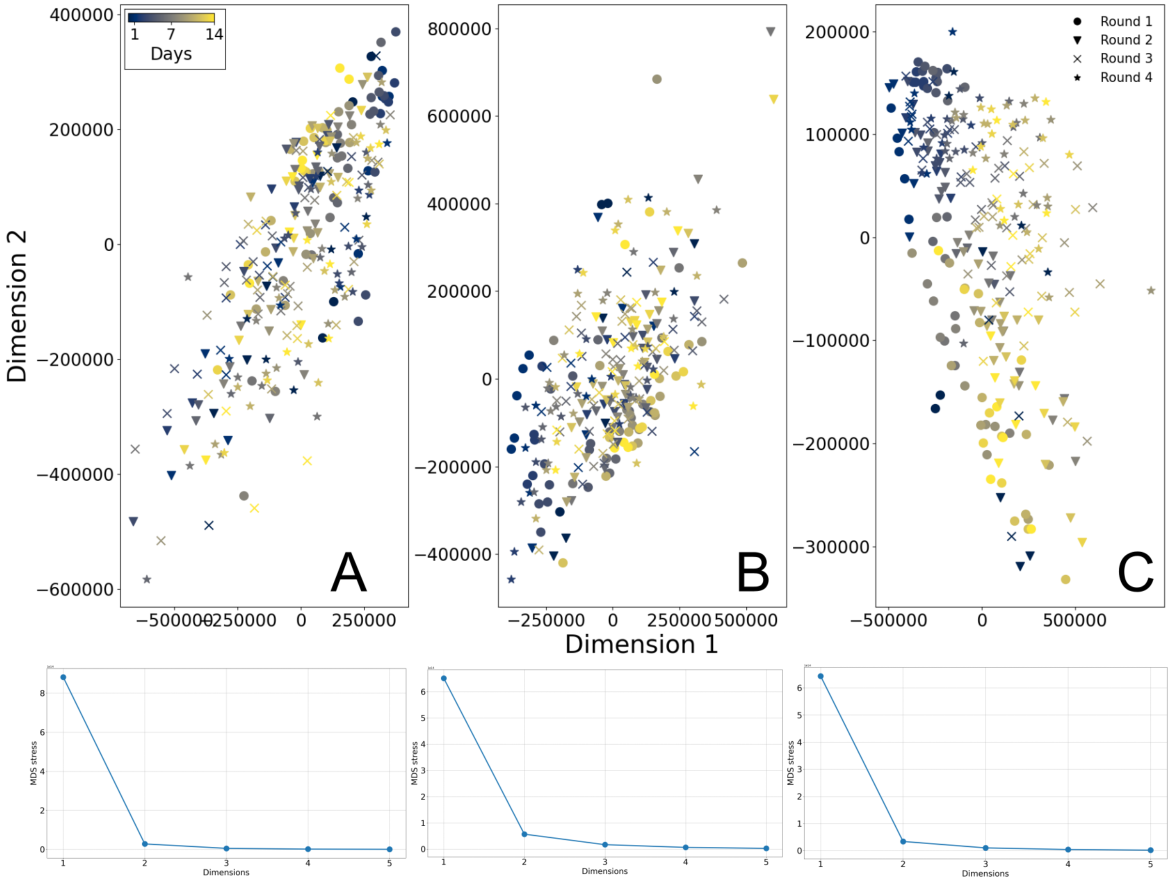


**Figure S24: Embedding Euclidean distances of EC number composition:** The Euclidean distances between communities were computed based on the EC number composition, as described in methods. The pairwise distances were embedded in two dimensions using multidimensional embedding for each environment **A** cellulose broth **B** cellulose paper and **C** glucose. The bottom panels show the stress of the embedding.

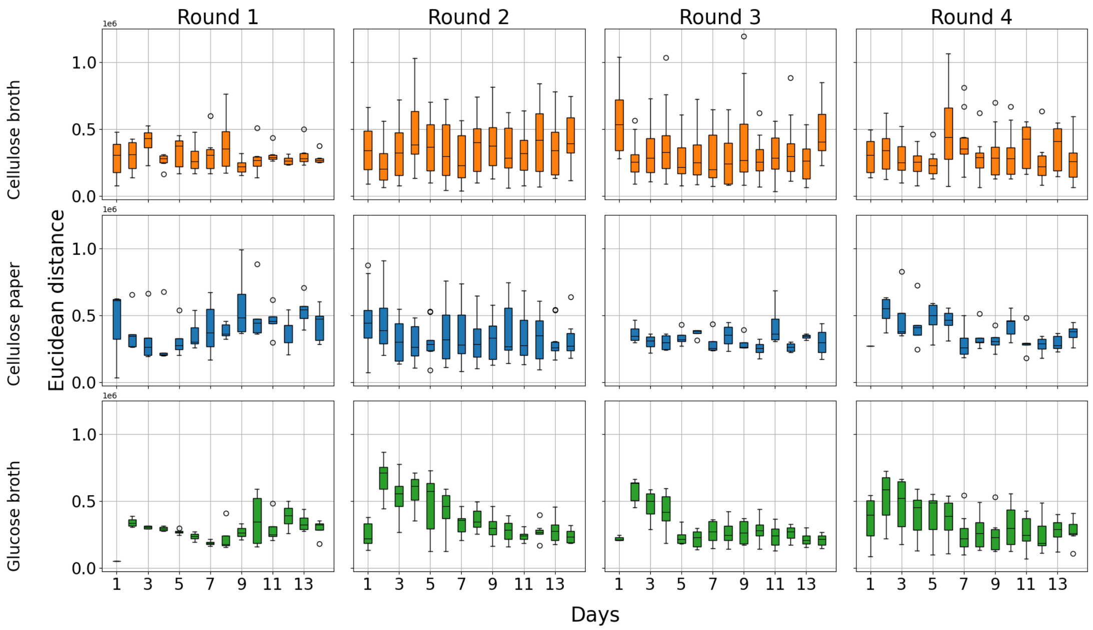


**Figure S25: Euclidean distances of EC composition across replicates:** The Euclidean distances between replicates based on EC composition were calculated. These are shown on the y axis. The x axis shows the days of sampling and the columns show the 4 rounds of transfer. The top row is for cellulose broth, the middle row for cellulose paper and bottom row for glucose. Each boxplot represents values across 5 replicate communities in each environment, where the boxes extend from the first to the third quartile, solid line represents the median, and the whiskers extend from the box to the farthest data point lying within 1.5 times the inter-quartile range.

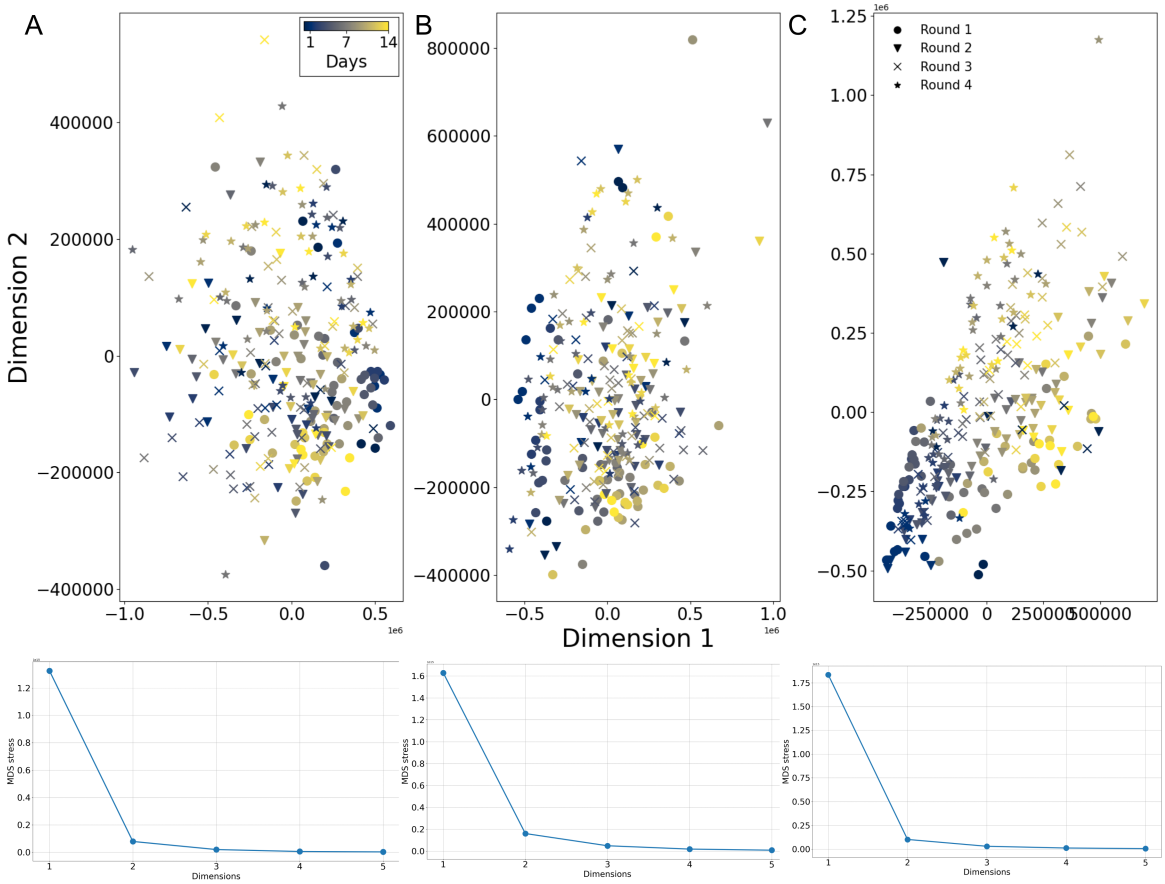


**Figure S26: Embedding Euclidean distances of KO composition:** The Euclidean distances between communities were computed based on the KO composition, as described in methods. The pairwise distances were embedded in two dimensions using multidimensional embedding for each environment **A** cellulose broth **B** cellulose paper and **C** glucose. The bottom panels show the stress of the embedding.

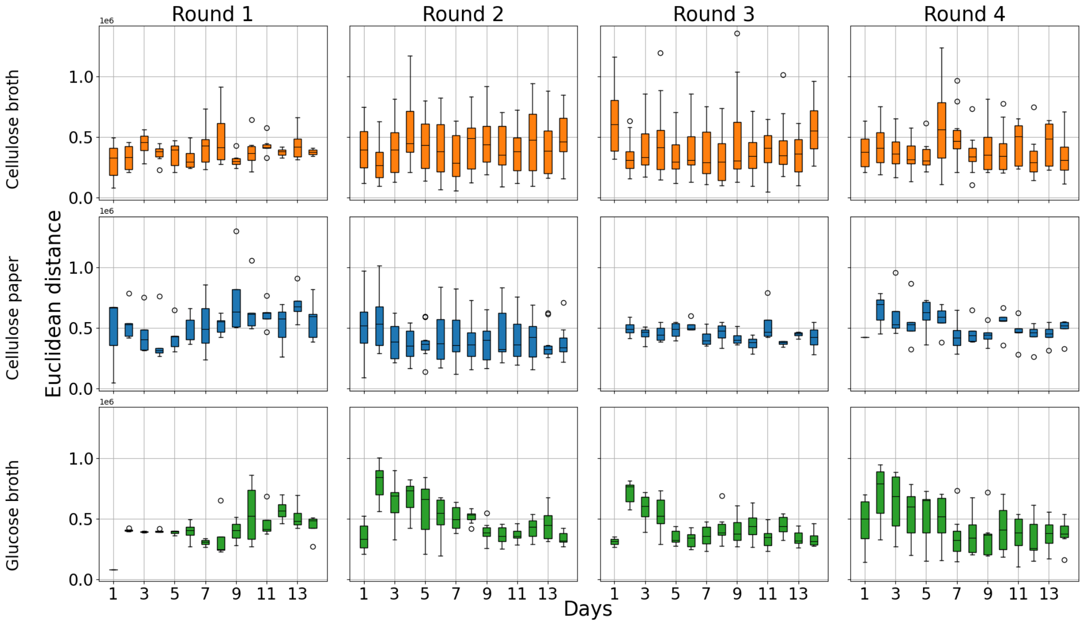


**Figure S27: Euclidean distances of KO composition across replicates:** The Euclidean distances between replicates based on KO composition were calculated. These are shown on the y axis. The x axis shows the days of sampling and the columns show the 4 rounds of transfer. The top row is for cellulose broth, the middle row for cellulose paper and bottom row for glucose. Each boxplot represents values across 5 replicate communities in each environment, where the boxes extend from the first to the third quartile, solid line represents the median, and the whiskers extend from the box to the farthest data point lying within 1.5 times the inter-quartile range.

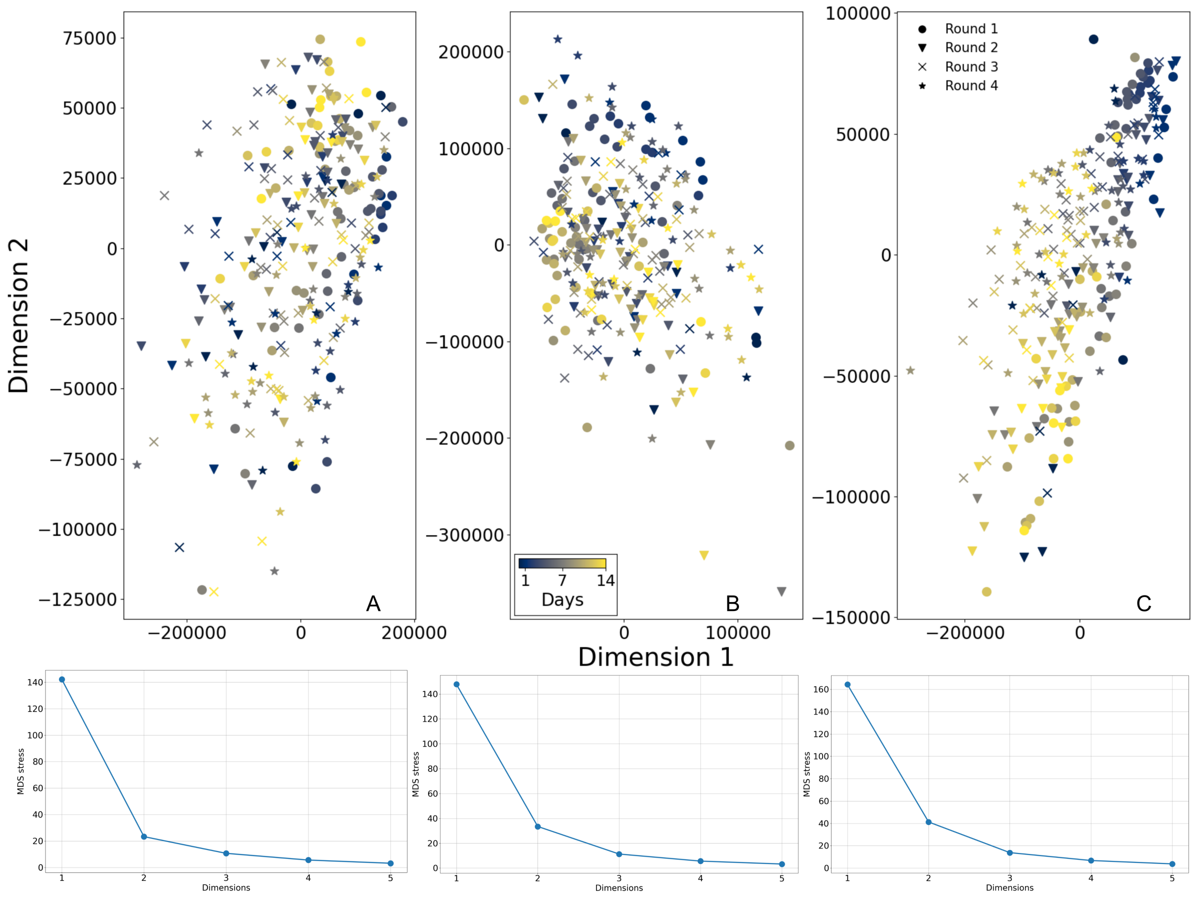


**Figure S28: Embedding Euclidean distances of pathways composition:** The Euclidean distances between communities were computed based on the pathways composition, as described in methods. The pairwise distances were embedded in two dimensions using multidimensional embedding for each environment **A** cellulose broth **B** cellulose paper and **C** glucose. The bottom panels show the stress of the embedding.

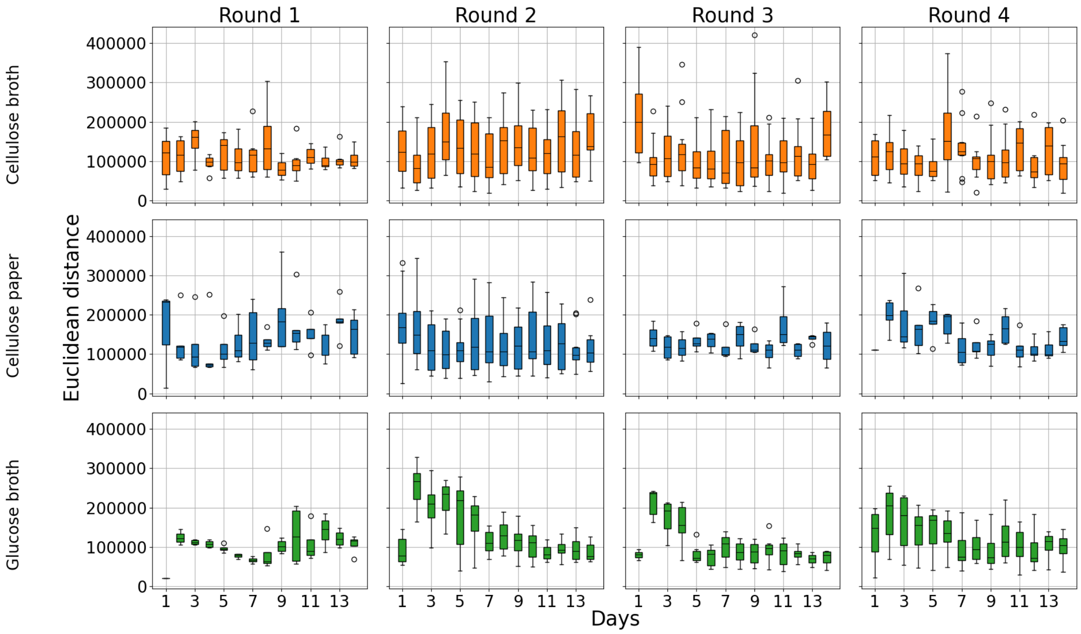


**Figure S29: Euclidean distances of pathways composition across replicates:** The Euclidean distances between replicates based on pathways composition were calculated. These are shown on the y axis. The x axis shows the days of sampling and the columns show the 4 rounds of transfer. The top row is for cellulose broth, the middle row for cellulose paper and bottom row for glucose.Each boxplot represents values across 5 replicate communities in each environment, where the boxes extend from the first to the third quartile, solid line represents the median, and the whiskers extend from the box to the farthest data point lying within 1.5 times the inter-quartile range.

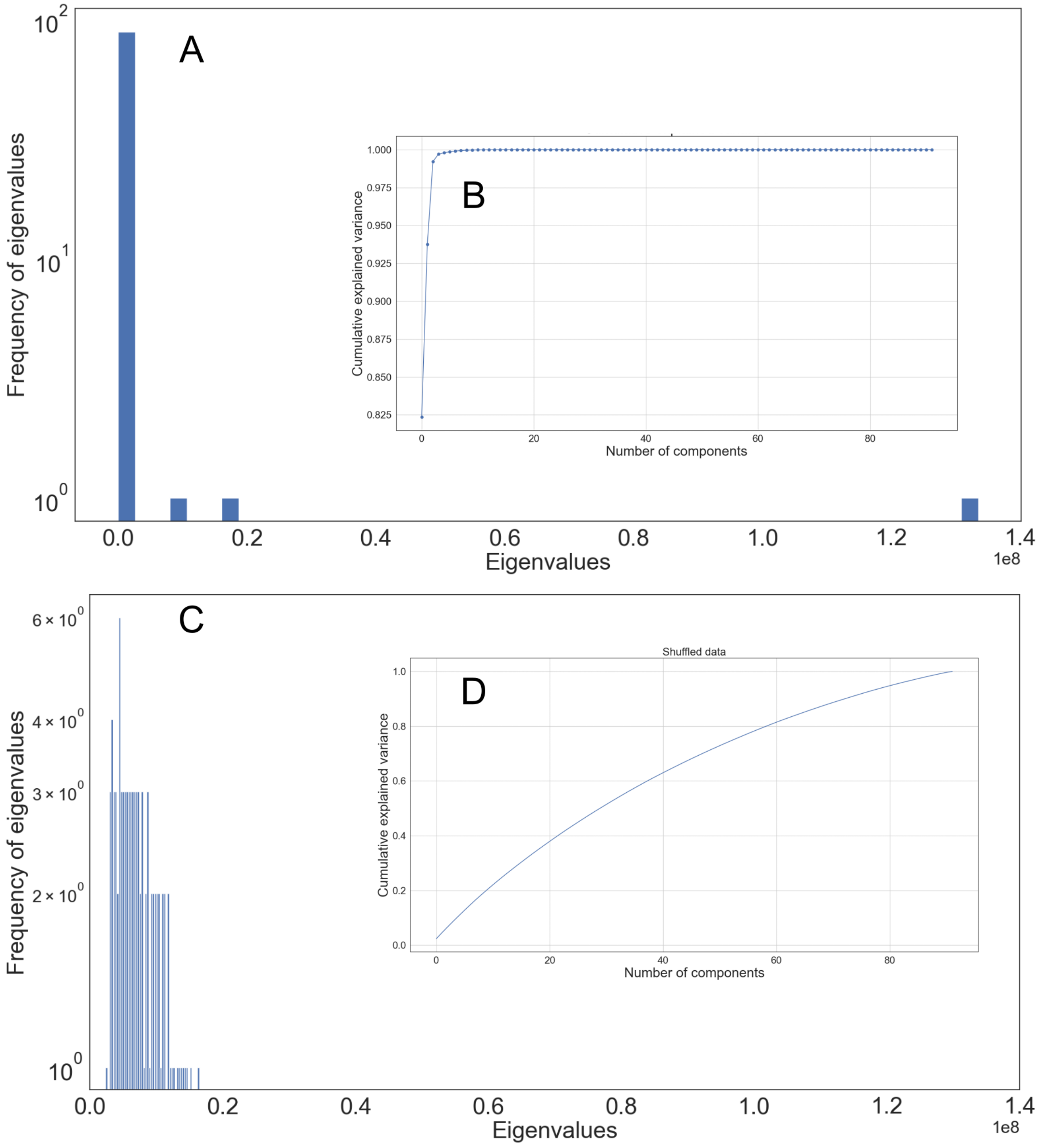


**Figure S30: Eigenvalues PCA on FAPROTAX output: A** shows the eigenvalues distribution of the FAPROTAX output **B** shows the cumulative explained variance. **C-D** show the eigenvalue distribution and cumulative explained variance for the shuffled data.

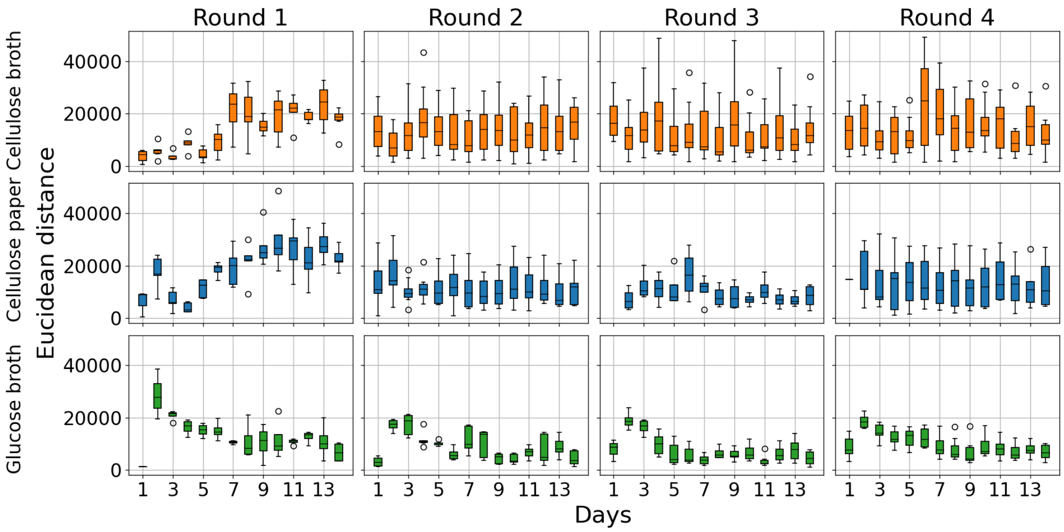


**Figure S31: Euclidean distances of functional composition across replicates:** The Euclidean distances between replicates based on the functional composition output of FAPROTAX were calculated. These are shown on the y axis. The x axis shows the days of sampling and the columns show the 4 rounds of transfer. The top row is for cellulose broth, the middle row for cellulose paper and bottom row for glucose.

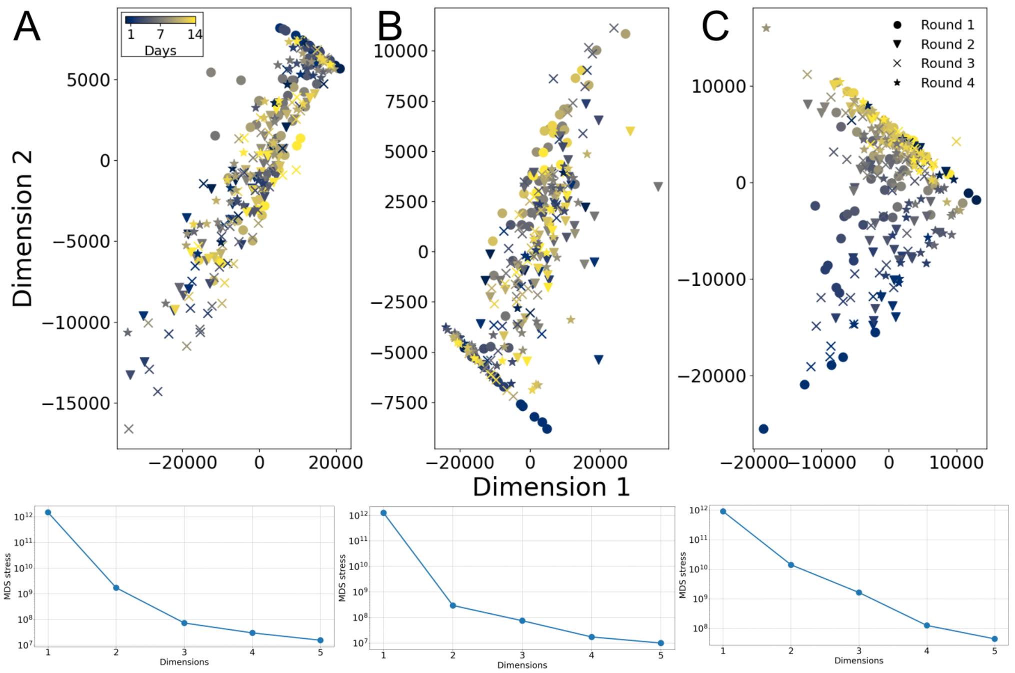


**Figure S32: Embedding Euclidean distances of functional composition provided by FAPROTAX:** The Euclidean distances between communities were computed based on the functional composition output of FAPROTAX, as described in methods. The pairwise distances were embedded in two dimensions using multidimensional embedding for each environment **A** cellulose broth **B** cellulose paper and **C** glucose. The bottom panels show the stress of the embedding. Each boxplot represents values across 5 replicate communities in each environment, where the boxes extend from the first to the third quartile, solid line represents the median, and the whiskers extend from the box to the farthest data point lying within 1.5 times the inter-quartile range.

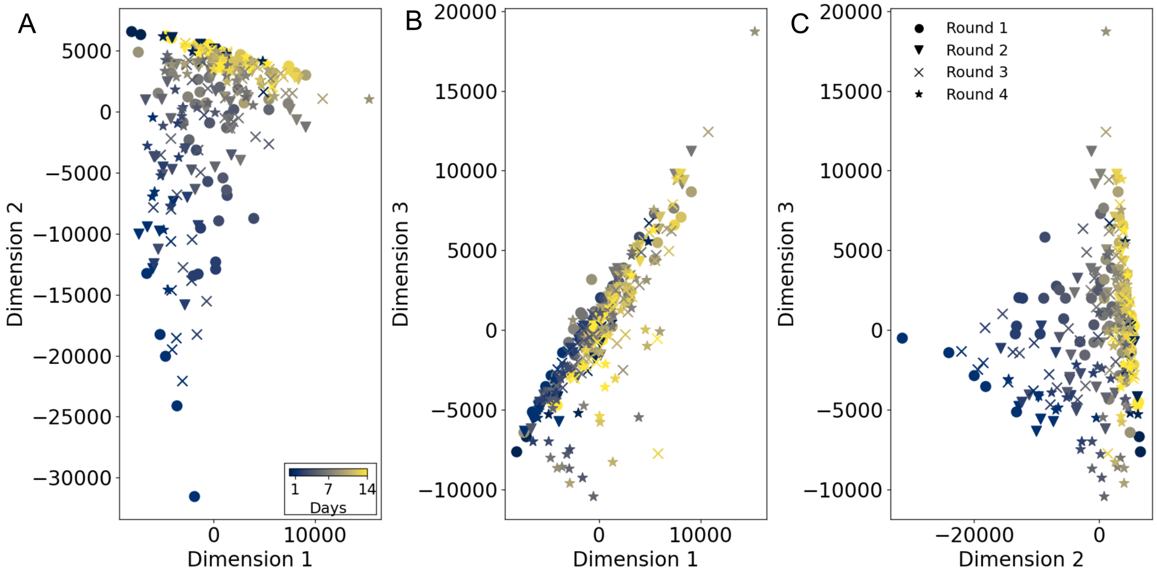


**Figure S33: 3 dimensional embedding of euclidean distances of functional composition provided by FAPROTAX of glucose communities:** The glucose communities were embedded in 3 dimensions to look for any additional clustering of the samples. **A-C** show the embeddings in Dimensions 1 and 2, 1 and 3 and 2 and 3 respectively.

| High in cellulose broth | High in glucose |
|---|---|
| DNA-directed DNA polymerase | glutathione transferase |
| DNA helicase | ABC-type polar-amino-acid transporter |
| NADH:ubiquinone reductase | sarcosine oxidase (formaldehyde-forming) |
| cytochrome-c oxidase | formate dehydrogenase |
| type I site-specific deoxyribonuclease | xanthine dehydrogenase |
| beta-glucosidase | taurine dioxygenase |
| methylmalonyl-CoA mutase | acetyl-CoA C-acetyltransferase |
| peptidylprolyl isomerase | enoyl-CoA hydratase |
| site-specific DNA-methyl-transferase (adenine-specific) | glutamine synthetase |
| cytochrome-c peroxidase | catalase |
| branched-chain-amino-acid transaminase | amidase |
| thioredoxin-disulfide reductase | urease |
| cysteine desulfurase | hippurate hydrolase |
| alpha-L-fucosidase | phosphoglycolate phosphatase |
| alanine dehydrogenase | nitronate monooxygenase |

**Table S1: Enzymes that have significantly higher abundances in the two environments.** Derived from 16S data using PICRUSt2, the enzymes whose abundance was higher in CB compared to glucose are listed in the first column, and the enzymes whose abundance was higher in the glucose-supported environment compared to the CB environment are listed in the second column.

| High in cellulose broth | High in glucose |
|---|---|
| putative ABC transport system | polar amino acid transport system substrate-binding protein |
| ABC-2 type transport system ATP-binding protein | polar amino acid transport system permease protein |
| LacI family transcriptional regulator | methyl-accepting chemotaxis protein |
| RNA polymerase sigma-70 factor (rpoE) | glutathione S-transferase |
| two-component system, LytTR family, response regulator | putrescine transport system substrate-binding protein |
| ATP-binding cassette | acyl-ACP dehydrogenase |
| ABC-2 type transport system permease protein | transmembrane sensor |
| putative ABC transport system ATP-binding protein | polar amino acid transport system ATP-binding protein |
| hydrophobic/amphiphilic exporter-1 | putative spermidine/putrescine transport system substrate-binding protein |
| cytochrome c oxidase subunit III | glycine cleavage system transcriptional activator |
| beta-glucosidase | glycine betaine/proline transport system substrate-binding protein |
| cytochrome c peroxidase | outer-membrane receptor for ferric coprogen and ferric-rhodotorulic acid |
| branched-chain amino acid aminotransferase | type VI secretion system secreted protein |
| thioredoxin reductase (NADPH) | D-methionine transport system substrate-binding protein |
| alanine dehydrogenase | formate dehydrogenase major subunit |

**Table S2: Kegg-orthologs (KOs) that have significantly higher abundances in the two environments.** Derived from 16S data using PICRUSt2, the KOs whose abundance was higher in CB compared to glucose are listed in the first column, and the KOs whose abundance was higher in the glucose-supported environment compared to the CB environment are listed in the second column.

| High in cellulose broth | High in glucose |
|---|---|
| L-valine biosynthesis | TCA cycle VII |
| L-isoleucine biosynthesis I | TCA cycle VI |
| L-isoleucine biosynthesis II | ornithine degradation |
| pyruvate fermentation to propionate | L-isoleucine biosynthesis IV |
| aerobic respiration I (cytochrome c) | stearate biosynthesis II |
| dTDP-N-acetylthomosamine biosynthesis | pyridoxal 5'-phosphate biosynthesis and salvage |
| superpathway of pyrimidine ribonucleosides salvage | palmitoleate biosynthesis I |
| Branched chain amino acid synthesis | L-aspartate and L-aspargine biosynthesis |
| pyrimidine deoxyribonucleosides salvage | protocatechuate degradation II |
| D-galacturonate degradation I | mycolate biosynthesis |
| adenine and adenosine salvage III | oleate biosynthesis IV |
| NAD de novo biosynthesis II (from tryptophan) | (5Z)-dodecenoate biosynthesis I |
| GDP-mannose-derived O-antigen building blocks biosynthesis | palmitate biosynthesis II |
| taxadiene biosynthesis (engineered) | fatty acid salvage |
| L-isoleucine biosynthesis III | adenosylcobalamin biosynthesis from adenosylcobinamide-GDP I |

**Table S3: Pathways that have significantly higher abundances in the two environments.** Derived from 16S data using PICRUSt2, the pathways whose abundance was higher in CB compared to glucose are listed in the first column, and the pathways whose abundance was higher in the glucose-supported environment compared to the CB environment are listed in the second column.

| High in cellulose broth | High in glucose |
|---|---|
| cellulolysis | nitrate reduction |
| aerobic chemoheterotrophy | fermentation |
| chemoheterotrophy | nitrogen respiration |
| ureolysis | nitrate respiration |
| phototrophy | human associated |
| photoheterotrophy | human pathogens all |
| | sulfate respiration |
| | respiration of sulfur compounds |
| | animal parasites or symbionts |

**Table S4: Functions that have significantly higher abundances in the two environments.** Derived from 16S data using FAPROTAX, the functions whose abundance was higher in CB compared to glucose are listed in the first column, and the functions whose abundance was higher in the glucose-supported environment compared to the CB environment are listed in the second column.

| High in the initial days | High in the final days |
|---|---|
| 6-phospho-beta-glucosidase | DNA-directed DNA polymerase |
| formate C-acetyltransferase | NADH:ubiquinone reductase (H+-translocating) |
| [formate-C-acetyltransferase] -activating enzyme | DNA 5'-3' helicase |
| pyruvate synthase | histidine kinase |
| glycerol dehydrogenase | peptidylprolyl isomerase |
| PepB aminopeptidase | cytochrome-c oxidase |
| acyl-Kdo2-lipid IVA acyltransferase | glutathione transferase |
| malate dehydrogenase (oxaloacetate-decarboxylating) | 3-oxoacyl-[acyl -carrier-protein] reductase |
| sugar-phosphatase | Glutamine synthetase |
| hydrogenase (acceptor) | acetyl-CoA C-acetyltransferase |

**Table S5: Enzymes that have significantly higher abundances in the early and later growth phases.** Derived from 16S data using PICRUSt2, the enzymes whose abundance was higher in the early growth phase compared to the later growth phase are listed in the first column, and those whose abundance was higher in the later growth phase compared to the early growth phase are listed in the second column.

| High in the initial days | High in the final days |
|---|---|
| 6-phospho-beta-glucosidase | RNA polymerase sigma-70 factor |
| serine transporter | iron complex outermembrane receptor protein |
| two-component system, LytTR family, sensor kinase | methyl-accepting chemotaxis protein |
| formate C-acetyltransferase | uncharacterized protein |
| pyruvate-ferredoxin/ flavodoxin oxidoreductase | ATP-binding cassette, subfamily B |
| concentrative nucleoside transporter, CNT family | glutathione S-transferase |
| putative transport protein | 3-oxoacyl-[acyl-carrier protein] reductase |
| MFS transporter, OPA family, sugar phosphate sensor protein UhpC | glutamine synthetase |
| glycerol dehydrogenase | acetyl-CoA C-acetyltransferase |
| C4-dicarboxylate transporter, DcuC family | putative ABC transport system permease protein |

**Table S6: KOs that have significantly higher abundances in the early and later growth phases.** Derived from 16S data using PICRUSt2, the KOs whose abundance was higher in the early growth phase compared to the later growth phase are listed in the first column, and those whose abundance was higher in the later growth phase compared to the early growth phase are listed in the second column.

| High in the initial days | High in the final days |
|---|---|
| (Kdo)2-lipid A biosynthesis | aerobic respiration I (cytochrome c) |
| thiazole biosynthesis II (aerobic bacteria) | pyruvate fermentation to isobutanol (engineered) |
| menaquinol-6 biosynthesis I | cis-vaccenate biosynthesis |
| menaquinol-9 biosynthesis | gondoate biosynthesis (anaerobic) |
| menaquinol-10 biosynthesis | L-isoleucine biosynthesis II |
| L-arginine and L-ornithine degradation | Valine synthesis |
| L-arginine, putrescine, and 4-aminobutanoate degradation | Isoleucine synthesis |
| heme b biosynthesis from uroporphyrinogen-III | fatty acid elongation – saturated |
| polymyxin resistance | mycolate biosynthesis |
| chorismate metabolism | oleate biosynthesis IV (anaerobic) |

**Table S7: Pathways that have significantly higher abundances in the early and later growth phases.** Derived from 16S data using PICRUSt2, the pathways whose abundance was higher in the early growth phase compared to the later growth phase are listed in the first column, and those whose abundance was higher in the later growth phase compared to the early growth phase are listed in the second column.

| High in the initial days | High in the final days |
|---|---|
| fermentation | aerobic chemoheterotrophy |
| nitrate reduction | nitrate respiration |
| | nitrogen respiration |
| | human pathogens all |
| | animal parasites or symbionts |
| | human associated |
| | sulfate respiration |
| | respiration of sulfur compounds |
| | intracellular parasites |
| | ureolysis |

**Table S8: Functions that have significantly higher abundances in the early and later growth phases.** Derived from 16S data using FAPROTAX, the functions whose abundance was higher in the early growth phase compared to the later growth phase are listed in the first column, and those whose abundance was higher in the later growth phase compared to the early growth phase are listed in the second column.
